# Loss of plastid developmental genes coincides with a reversion to monoplastidy in hornworts

**DOI:** 10.1101/2022.01.11.475830

**Authors:** Alexander I. MacLeod, Parth K. Raval, Simon Stockhorst, Michael R. Knopp, Eftychios Frangedakis, Sven B. Gould

## Abstract

The first plastid evolved from an endosymbiotic cyanobacterium in the common ancestor of the Archaeplastida. The transformative steps from cyanobacterium to organelle included the transfer of control over developmental processes; a necessity for the host to orchestrate, for example, the fission of the organelle. The plastids of almost all embryophytes divide independent from nuclear division, leading to cells housing multiple plastids. Hornworts, however, are monoplastidic (or near-monoplastidic) and their photosynthetic organelles are a curious exception among embryophytes for reasons such as the occasional presence of pyrenoids. Here we screened genomic and transcriptomic data of eleven hornworts for components of plastid developmental pathways. We find intriguing differences among hornworts and specifically highlight that pathway components involved in regulating plastid development and biogenesis were differentially lost in this group of bryophytes. In combination with ancestral state reconstruction, our data suggest that hornworts have reverted back to a monoplastidic phenotype due to the combined loss of two plastid division-associated genes: ARC3 and FtsZ2.

## INTRODUCTION

Hornworts are a unique group of bryophytes, the monophyletic non-vascular sister lineage to all vascular land plants (Harris et al. 2020). The phylogenetic position of hornworts, and their putative phenotypic resemblance to what one might consider to represent the last common ancestor of all land plants, make them an attractive model for evo-devo studies linked to events such as plant terrestrialization (Frangedakis et al. 2020). Hornworts are the only group of land plants known to form a pyrenoid, a unique carbon concentrating mechanism (CCM) otherwise common in algae – however, these CCMs are not present in all hornworts and are hence a poor taxonomic marker (*Figure 1*) (Villarreal & Renner 2012).

**Figure 1.**
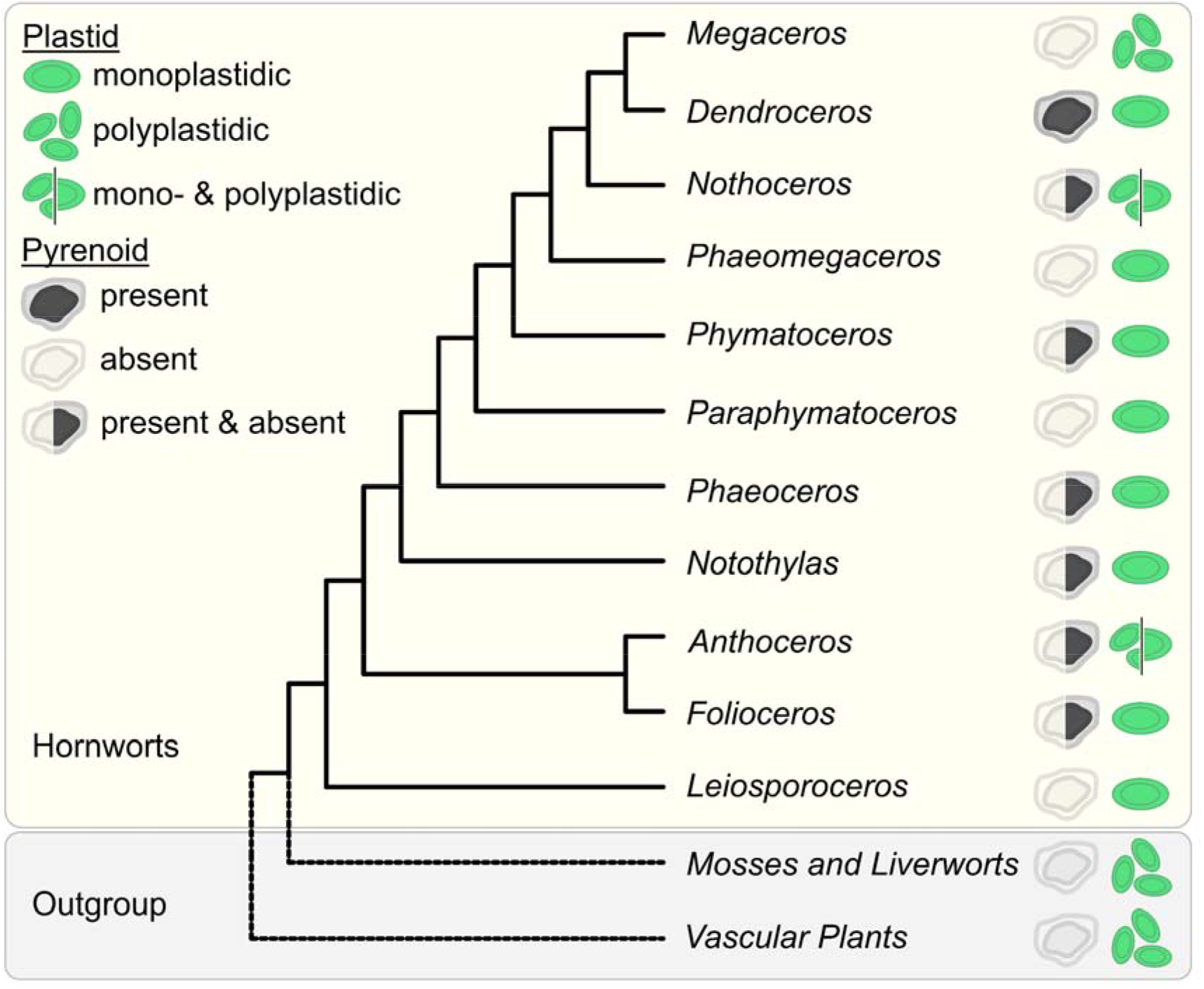
Maximum likelihood (ML) phylogeny of the main eleven hornwort families. Pyrenoidal and thallus/gametophyte plastidic phenotypes for the genera are indicated, based on Vaughan *et al*. (1992), Li *et al*. (2017), Villarreal *et al*. (2012) and Raven & Edwards (2014). Outgroups are highlighted by the dotted lines.

Hornworts are one of the only groups of embryophytes that have not escaped the monoplastidic bottleneck. It is a phenomenon associated with plastid origin and the organelle’s integration into the host cell cycle, which constrain the majority of algae from possessing multiple plastids per cell (*Figure 1*) (de Vries & Gould 2018). One consequence is that the only plastids that hornwort cells house are chloroplasts, whose size and morphology vary across genera (Raven & Edwards 2014; Vaughn et al. 1992; Li et al. 2017). To address why, we screened the genomes and annotated transcriptomes of ten hornwort species to identify the presence/absence of genes that play key roles in regulating plastid development (Jarvis & López-Juez 2013). We highlight key differences between the developmental plastid biology of hornworts and other established model organisms in the terrestrial clade. Furthermore, we argue that major changes in plastid biology not only coincided with major checkpoints in the evolutionary history of hornworts, but also facilitated them.

## MATERIALS AND METHODS

### Species and gene tree phylogeny constructions

A maximum likelihood (ML) tree (*Figure 1*) was constructed via IQ-TREE version 2.0.3 (Minh et al. 2020), using an automated selection model, by concatenating single-copy chloroplast and mitochondrial markers from 65 different hornwort species, and three outgroups (Villarreal & Renner 2012). Said sequences were aligned with MUSCLE in AliView (Laarson 2014; Edgar 2004). Gene trees for orthologues were generated using PhyML version 3.0 and IQ-TREE version 2.0.3 using automated selection models (Guindon et al. 2010; Lefort et al. 2017). We used the SHOOT framework (Emms & Kelly 2021) to extract orthologous sequences from across the Archaeplastida.

### Orthologue classification and identification

We analysed the genomes and transcriptomes of ten hornworts (supplementary table S1), along with the genomes of *Arabidopsis thaliana* and *Marchantia polymorpha*, to determine the presence of various components involved in plastid development (Bowman et al. 2017; Lamesch et al. 2012; Leebens-Mack et al. 2019; Zhang et al. 2020; Li et al. 2020). To estimate the completeness of each annotation, we used BUSCO version 5.2.2 (supplementary table S1) (Manni et al. 2021). Orthology clusters (Orthogroups) were identified using OrthoFinder version 2.5.4 (Emms & Kelly 2019, 2015) (supplementary table S2). To validate Orthogroup presence/absence, we checked for reciprocal best hits using DIAMOND (Buchfink et al. 2015). Due to the difficulty in identifying orthologues for the import protein YCF1 in the Archaeplastida (de Vries et al. 2015), we employed a different strategy to identify orthologues for this gene. We extracted established YCF1 sequences from GenBank and UniProt and used them as queries for DIAMOND.

### Ancestral state reconstructions (ASRs)

A robust ML species phylogeny of the green lineage was constructed via IQ-TREE version 2.0.3 (Minh et al. 2020), using an automated selection model, by concatenating several housekeeping genes identified with DIAMOND (supplementary table S3 and S4) (Buchfink et al. 2015). We used a reciprocal best hit pipeline with DIAMOND (Buchfink et al. 2015), to analyze the genomes of 34 different Streptophytes, seven Chlorophytes and one Glaucophyte (supplementary table S3) to determine the presence and absence of orthologues involved in plastid division, to estimate the presence/absence of ARC3 and FtsZ2 at various nodes on our tree (supplementary table S5). Subsequent ASRs were undertaken using the ape function from the Phytools package (Revell 2012).

## RESULTS AND DISCUSSION

### Full conservation of TOC but only partial conservation of TIC in hornwort chloroplasts

The vast majority of plastid proteins are encoded by the nuclear genome and, after their synthesis in the cytosol, are imported into the plastid by the TOC/TIC (translocon of the outer/inner envelope of the chloroplast) complex (Richardson & Schnell 2020). Embryophytes have evolved the most sophisticated TOC/TIC complexes (Knopp et al., 2020) and our data confirm that the hornwort TOC complex is comprised of the same key proteins that are found in other embryophytes, mainly TOC75, TOC34 and TOC159 (Richardson & Schnell 2020) (*Figure 2A* and supplementary figures S7-19). The recycling of major TOC components is regulated by the RING-type ubiquitin E3 ligase SP1, which targets these proteins for proteasomal degradation (Ling et al. 2012) (*Figure 2B*).

**Figure 2.**
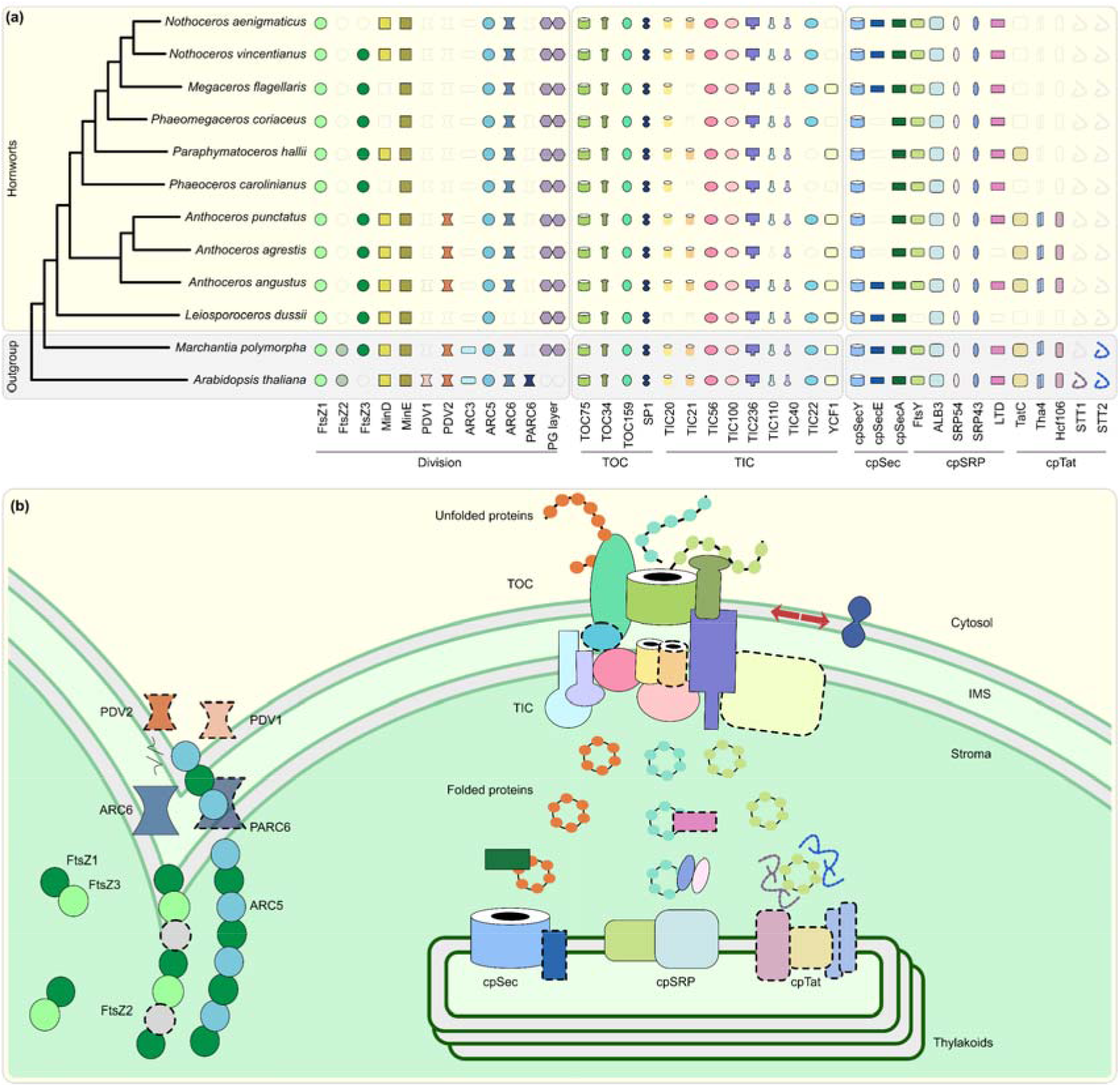
Plastid development and biogenesis in hornworts. **(a)** A presence/absence pattern (PAP) of various plastid developmental components that are sorted into three categories based on whether they are associated with plastid division (PD), protein translocation across the plastid envelope via TOC/TIC or the thylakoid membrane. Transparent icons indicate that no gene could be identified. **(b)** A combined schematic representation of plastid development in embryophytes. Components that are absent from more than two hornworts in our surveyed taxa, or absent in this group altogether, are highlighted by dotted outlines. ARC, accumulation and regulation of chloroplasts; FtsZ, filamentous temperature Z; IMS, intermembrane space; Sec, secretory; SRP, signal recognition particle; Tat; twin arginine translocation; TOC/TIC, translocator of the outer/inner chloroplast membrane; PDV, plastid division. While ARC5 is absent from the Anthoceros agrestis Bonn ecotype, which we included in our OrthoFinder analyses as the representative for this species, our reciprocal best hit pipeline confirmed that it is present in the Oxford ecotype, with its gene ID being AagrOXF_evm.TU.utg000081l.174.

The TIC complex of embryophytes is comprised of a 1 MDa multimer that forms a pore that receives precursor protein from the TOC complex in the intermembrane space (IMS), and finally mediates their passage to the stroma (Nakai 2015a; Richardson & Schnell 2020). The presence/absence of TOC/TIC components reveal no pattern with regard to mono-/polyplastidy or presence/absence of a pyrenoid (*Figure 2A, Figure 1*). However, some TIC components appear to have undergone differential loss in some hornwort taxa (*Figure 2A*), most notably TIC21, TIC22, YCF1 (TIC214), and maybe even TIC20 in *Leiosporoceros dussii*. The latter species is the only member of our surveyed taxa that lacks a TIC20 orthologue (*Figure 2A*). This could be the result of a transcriptome annotation and coverage issues (Cheon et al. 2020), since TIC20 is hypothesized to be a universal protein across the green lineage (Kalanon & McFadden 2008; de Vries et al. 2015). Should this not be the case, then maybe YCF1/TIC214 and TIC100 can compensate for TIC20’s absence in a unique manner.

YCF1/TIC214, the only TOC/TIC component encoded by the plastid genome and unique to the green lineage, is absent from a significant number of hornworts (*Figure 2A*) such as in *Nothoceros aenigmaticus* for which also the plastid genome is available (Villarreal et al. 2013). Some putative absences of YCF1/TIC214 could be the result of assembly and/or annotation errors, however, the gene was lost without question in grasses, too (de Vries et al. 2015; Nakai 2015b). The loss of this import protein does not necessarily lead to the loss of the entire import capacity (Bölter & Soll 2017) and raises the question whether there is a functional, causative correlation between the loss of YCF1/TIC214 across these diverse embryophyte groups.

### Differential loss of an ancient thylakoid developmental pathway in most hornworts

Thylakoid proteomes contain the bulk of photosynthesis-related proteins of plant cells (Xu et al., 2021). After their import via TOC/TIC, thylakoid proteins are recognized and sorted via one of three main pathways, the components of which are predominantly derived from the cyanobacterial endosymbiont, or inserted spontaneously (*Figure 2*, supplementary figures S20-30) (Xu et al., 2021).

The chloroplast secretory (cpSec) pathway is involved in importing unfolded proteins to the thylakoid lumen. Powered by the motor protein cpSecA, unfolded subunits pass through a pore formed by cpSecY and cpSecE (Xu et al. 2021). Half of surveyed hornworts lack cpSecE orthologues, with this distribution not showing any unique phylogenetic pattern (*Figure 1, Figure 2A*). Considering cpSecE only plays an accessory role in protein translocation by tiling and rotating cpSecY’s N-terminal half, its absence in some hornworts indicate that it might not be detrimental to the function of the cpSec pathway (Park et al. 2014).

The chloroplast twin-arginine translocation (cpTat) pathway can import folded proteins and is powered by the thylakoid’s proton motive force (PMF) (Xu et al. 2021). In those hornworts, for which we identified the cpTat pathway, it is comprised of three proteins: Tha4, TatC and Hcf106 (*Figure 2*). Precursor proteins initially bind to a TatC-Hcf106 complex. Tha4 is subsequently recruited via the action of the PMF, undergoing a conformational change, leading to the passage of the precursor protein (Xu et al. 2021) (*Figure 2B*). The cpTat pathway seems only to be encoded by the Anthocerotaceae, having been lost in other hornwort families (*Figure 2A*). If the cpTat pathway is indeed absent in most hornwort families, then this raises the question on how the thylakoids import folded proteins. Furthermore, all hornworts appear to lack STT proteins (*Figure 2A*), which mediate liquid-liquid phase transitions (LLPTs), allowing for more efficient sorting of cpTat substrates (*Figure 2A*) (Ouyang et al. 2020). cpTat-related LLPTs hence appear absent in hornworts or are regulated otherwise.

The third main pathway involved in sorting proteins for thylakoid biogenesis is the chloroplast signal recognition particle (cpSRP) pathway. This translocation complex is involved in targeting specifically light harvesting complex (LHCP) proteins to the thylakoid membrane (Xu et al. 2021) (*Figure 2B*). LHCP integration is initiated when a rudimentary LHCP is transferred from the TIC translocon to the SRP43/SRP54 complex by the LTD protein. Subsequently, this SRP43/SRP54 complex binds to the FtsY receptor. GTP hydrolysis results in LHCP integration via the action of the ALB3 integral translocase (Xu et al. 2021). Our results suggest that the cpSRP pathway is ubiquitous in all hornworts, as the core components of this pathway are present in the vast majority of our surveyed taxa. However, FtsY is absent in *L. dussii* and LTD is absent in both *Anthoceros angustus* and *L. dussii*. This differential loss of FtsY and LTD in *L. dussii* could be a consequence of this species potentially losing TIC20, with this core TIC component being a key LTD interaction partner (Ouyang et al. 2011).

### Loss of plastid division components coincide with monoplastidy in hornworts

Plastid division in bryophytes is achieved by three components: the outer and inner rings and most likely the peptidoglycan (PG) layer. The inner division ring (Z-ring) is comprised of FtsZ1,FtsZ2 and FtsZ3, while the outer division ring comprises ARC5 and FtsZ3 (Osteryoung & Pyke 2014; Grosche & Rensing 2017). Z-ring and outer ring synchronization are achieved via an interplay of ARC6 and PDV2 (Osteryoung & Pyke 2014). The PG layer is a relic of the chloroplast’s cyanobacterial past, and it might be relevant in regulating chloroplast division in bryophytes and streptophyte algae (Grosche & Rensing 2017; Hirano et al. 2016). We find that the chloroplasts of all surveyed hornworts possess all the enzymes necessary for PG layer biosynthesis (*Figure 2A* and *2B*), hinting towards a conserved function similar to that in the moss *Physcomitrium patens* (Hirano et al. 2016).

Hornworts appear to have differentially lost *both* ARC3 and FtsZ2 (*Figure 2*). This differential loss correlates with this group of bryophytes reverting back to a monoplastidic, or near-monoplastidic, phenotype (*Figure 1*) (Villarreal & Renner 2012; Li et al. 2017; Raven & Edwards 2014). Previous studies have shown that generating individual gene mutant lines of *ARC3* and *FtsZ2* in *A. thaliana* and *P. patens* cause fewer plastids (in the case of *arc3* mutants) or one giant plastid per cell (in the case of *ftsz2* mutants) (Pyke & Leech 1992; Martin et al. 2009). ARC3 is part of the FtsZ family and unites an FtsZ domain with a C-terminal MORN domain (Zhang et al. 2013). While ARC3 orthologues are absent in some polyplastidic seedless plants (such as *P. patens* and the lycophyte *Selaginella moellendorffii*), these species then possess orthologues for FtsZ2, which might compensate its loss to some degree (Albert et al. 2011; Rensing et al. 2008; Zhang et al. 2013). This is further supported by an ancestral state reconstruction analysis that demonstrates that the ancestral embryophyte possessed both ARC3 and FtsZ2 and was polyplastidic; the opposite of which is true for the ancestral hornwort (*Figure 3*, supplementary figures S31 and S32). We predict that the loss of both genes contributed to the monoplastidic nature of hornworts and that re-introducing them might induce a polyplastidic phenotype.

**Figure 3.**
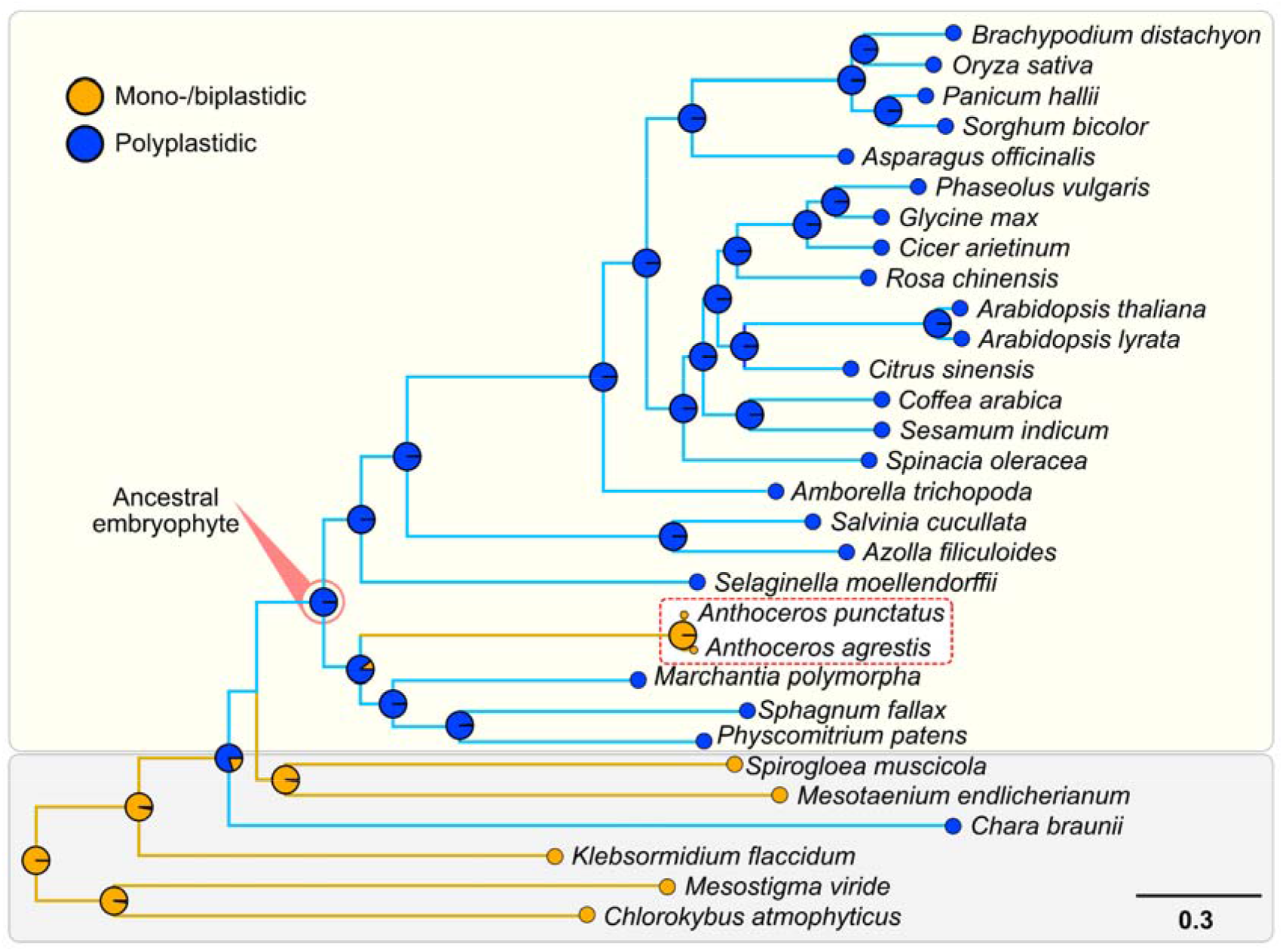
Support for the polyplastidic nature of the ancestral embryophyte and monoplastidic nature of the ancestral hornwort. Pie charts at the nodes display estimates of the probabilities for the plastidic phenotype of the respective most recent common ancestors (MRCAs). Hornworts are highlighted with a white box and a red dotted line.

## CONCLUDING REMARKS

It is evident that hornwort – and bryophyte – emergence and diversification was accompanied by major instances of gene loss (Harris et al. 2021). We suggest that a consequence of some of plastid-related gene losses, namely the combined loss of FtsZ2 and ARC3, resulted in hornworts reverting back to a monoplastidic phenotype, which the embryophyte ancestor was able to escape. If the knockout of ARC3 and FtsZ2 in *A. thaliana* and *P. patens* results in monoplastidic phenotypes, could one reverse evolution by expressing ARC3 and/or FtsZ2 in a hornwort? We anticipate our study to be a starting point for further experiments aimed at deconstructing bryophyte plastid biology and reconstructing new evolutionary hypotheses for outstanding questions in this topic. Next to exploring the monoplastidic bottleneck, hornworts might be able to shed new light on the import of folded proteins into the thylakoid of non-Anthocerotaceae hornworts, or the consequences of a potential TIC20 loss in *L. dussii* and the detailed function of YCF1, which next to grasses also some hornworts appear to have lost.

## DATA AVAILABILITY

All data generated in our study can be accessed at https://figshare.com/account/login#/projects/125431.

## ACKNOWLEDGEMNTS

We are grateful for the support by the DFG (SFB 1208–267205415 and SPP2237–440043394) and the VolkswagenStiftung (Life). AIM is furthermore supported by the Moore and Simons Initiative grant (9743) of William F. Martin.

