## Supplementary Figures S1-S32 for "Loss of plastid developmental genes coincides with a reversion to monoplastidy in hornworts"

### Contents

Supplementary Figures S1-S6 (Plastid Division) 1

Supplementary Figures S7-S19 (Protein Import to the Plastid) 7

Supplementary Figures S20-S30 (Thylakoid Development) 20

Ancestral State Reconstruction analyses for key PD Machinery Genes 31

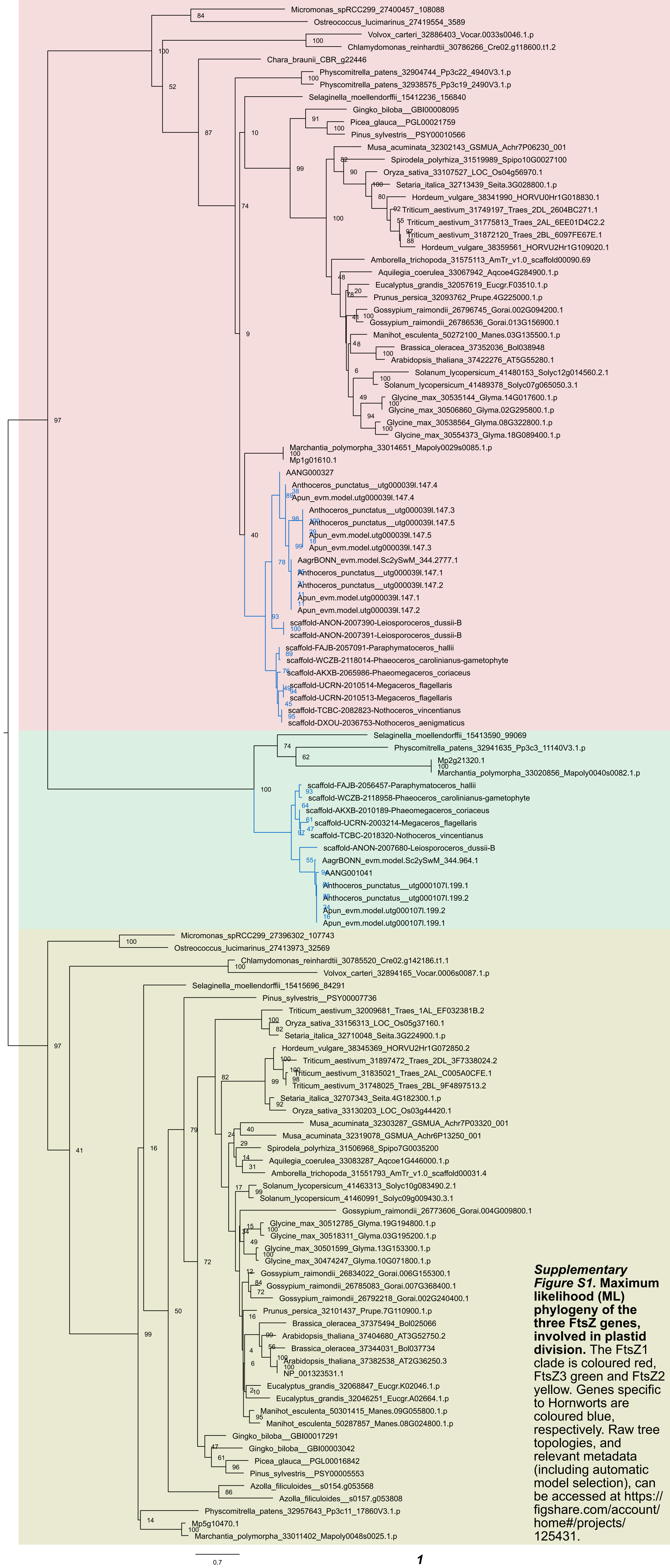


***Supplementary Figure S1.*** **Maximum likelihood (ML) phylogeny of the three FtsZ genes, involved in plastid division.** The FtsZ1 clade is coloured red, FtsZ3 green and FtsZ2 yellow. Genes specific to Hornworts are coloured blue, respectively. Raw tree topologies, and relevant metadata (including automatic model selection), can be accessed at https://figshare.com/account/home#/projects/125431.

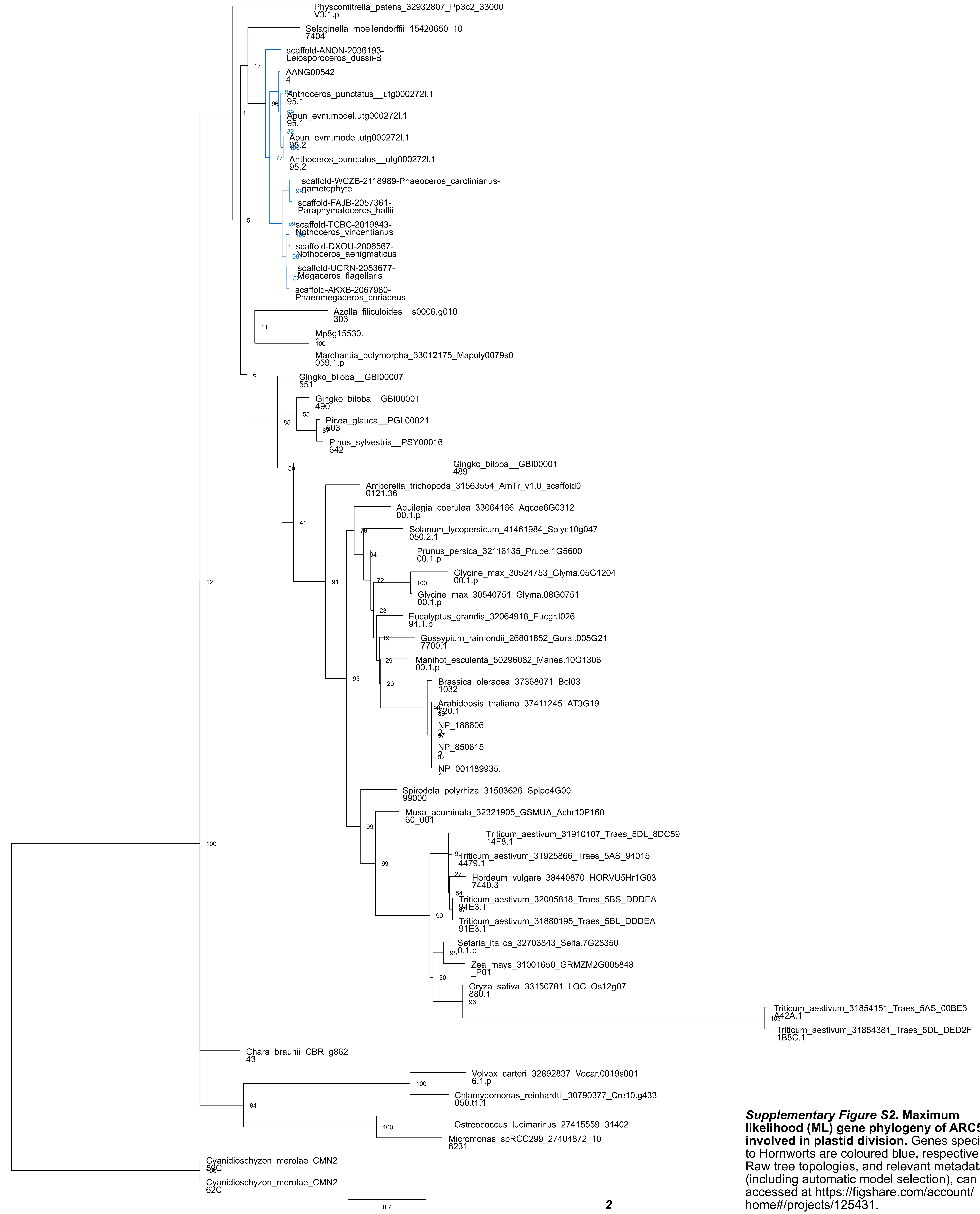


***Supplementary Figure S2.* Maximum likelihood (ML) gene phylogeny of ARC5, involved in plastid division.** Genes specific to Hornworts are coloured blue, respectively. Raw tree topologies, and relevant metadata (including automatic model selection), can be accessed at https://figshare.com/account/home#/projects/125431.

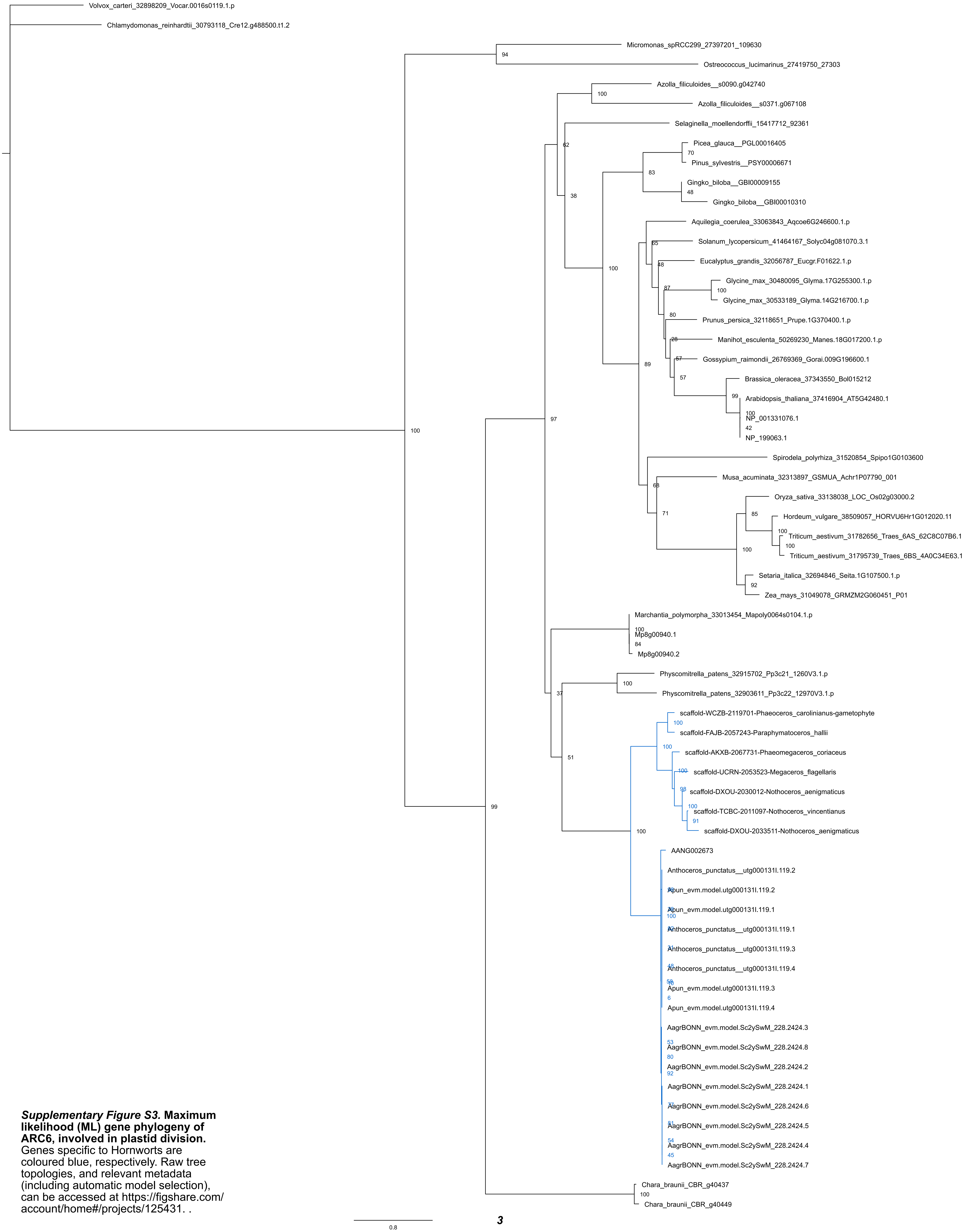


***Supplementary Figure S3.*** **Maximum likelihood (ML) gene phylogeny of ARC6, involved in plastid division.** Genes specific to Hornworts are coloured blue, respectively. Raw tree topologies, and relevant metadata (including automatic model selection), can be accessed at https://figshare.com/account/home#/projects/125431. .

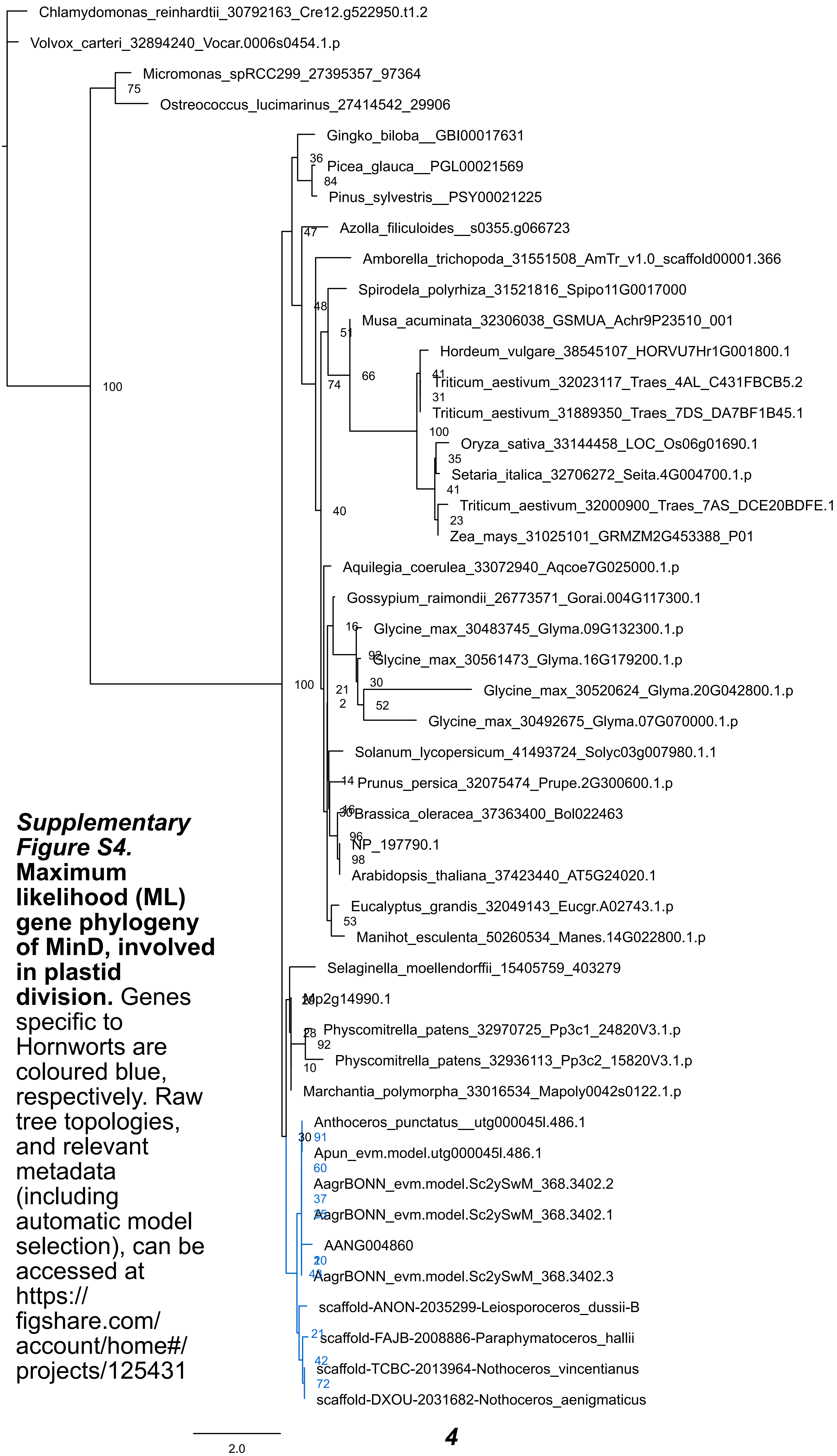


***Supplementary Figure S4.* Maximum likelihood (ML) gene phylogeny of MinD, involved in plastid division.** Genes specific to Hornworts are coloured blue, respectively. Raw tree topologies, and relevant metadata (including automatic model selection), can be accessed at https://figshare.com/account/home#/projects/125431

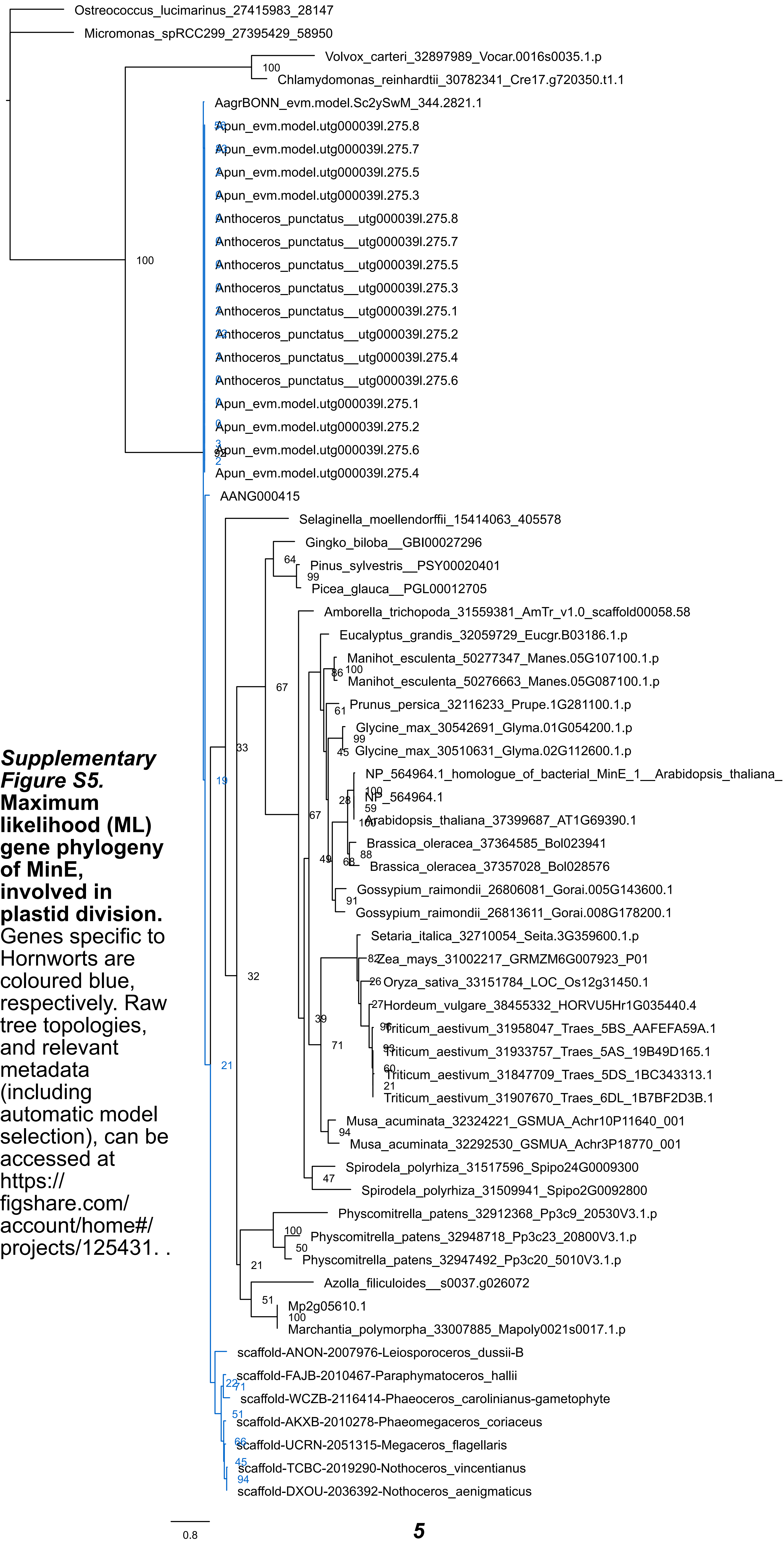


***Supplementary Figure S5.*** **Maximum likelihood (ML) gene phylogeny of MinE, involved in plastid division.** Genes specific to Hornworts are coloured blue, respectively. Raw tree topologies, and relevant metadata (including automatic model selection), can be accessed at https://figshare.com/account/home#/projects/125431. .

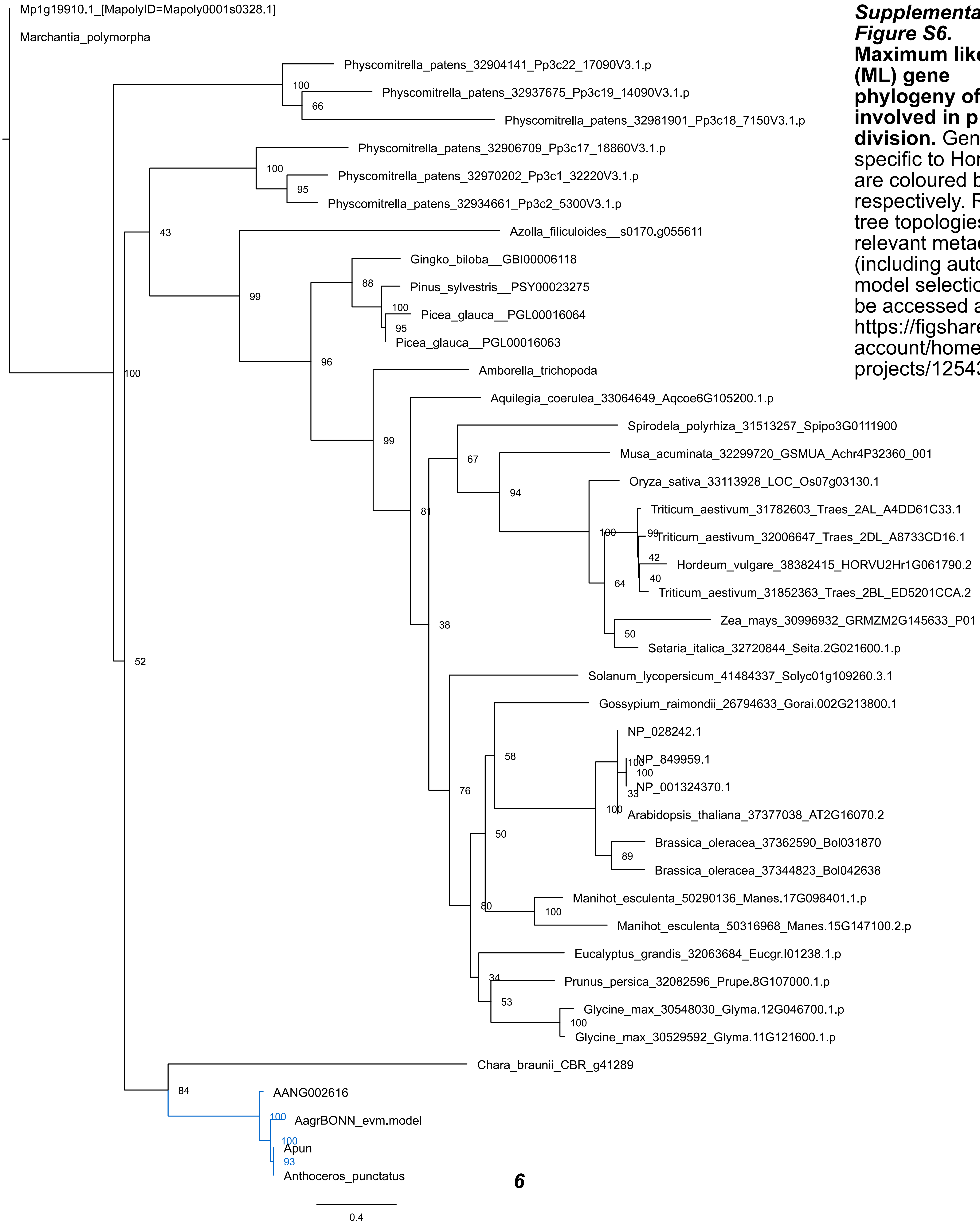


***Supplementary Figure S6.* Maximum likelihood (ML) gene phylogeny of PDV2, involved in plastid division.** Genes specific to Hornworts are coloured blue, respectively. Raw tree topologies, and relevant metadata (including automatic model selection), can be accessed at https://figshare.com/account/home#/projects/125431.

*6*

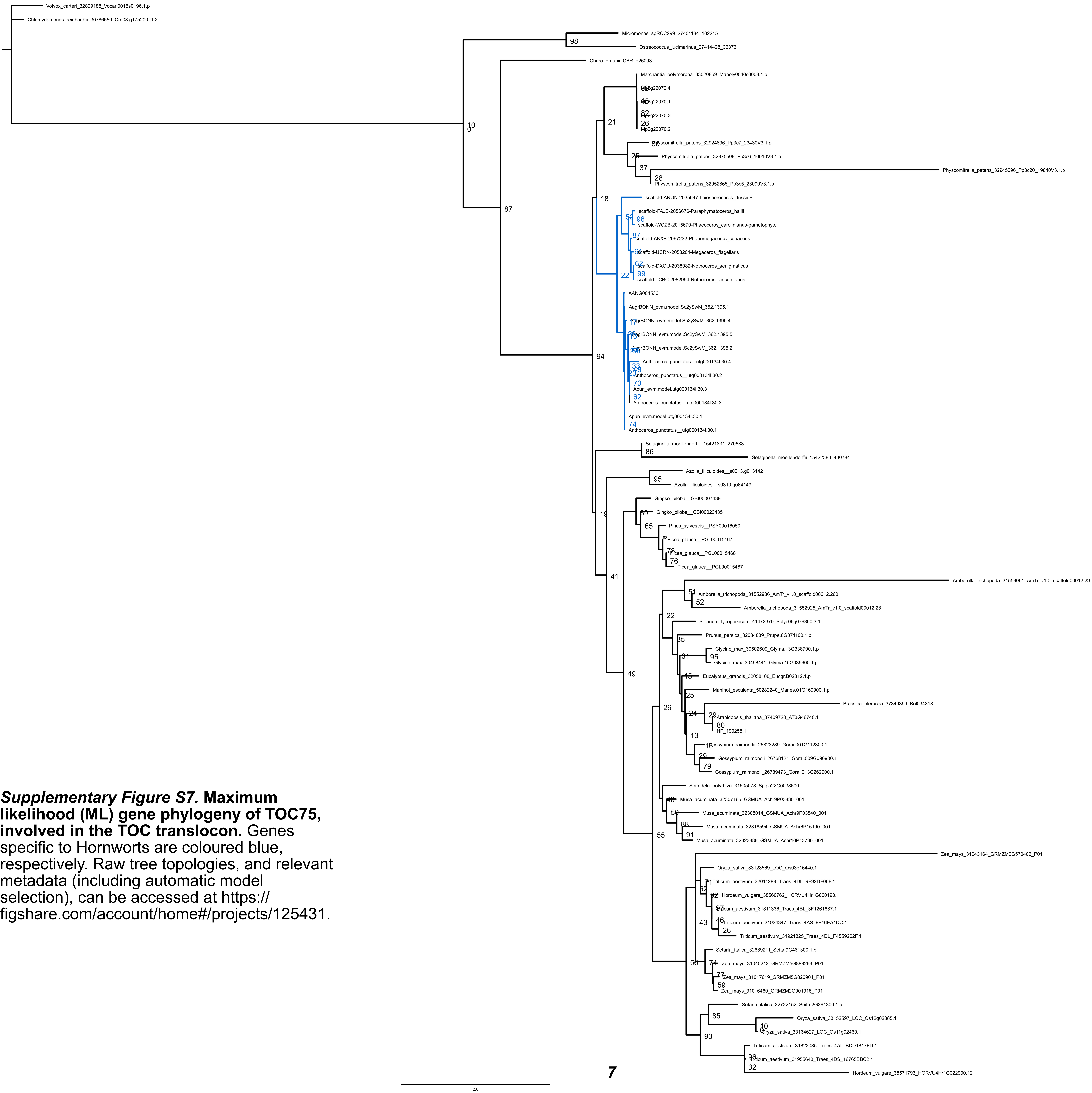


***Supplementary Figure S7.* Maximum likelihood (ML) gene phylogeny of TOC75, involved in the TOC translocon.** Genes specific to Hornworts are coloured blue, respectively. Raw tree topologies, and relevant metadata (including automatic model selection), can be accessed at https://figshare.com/account/home#/projects/125431.

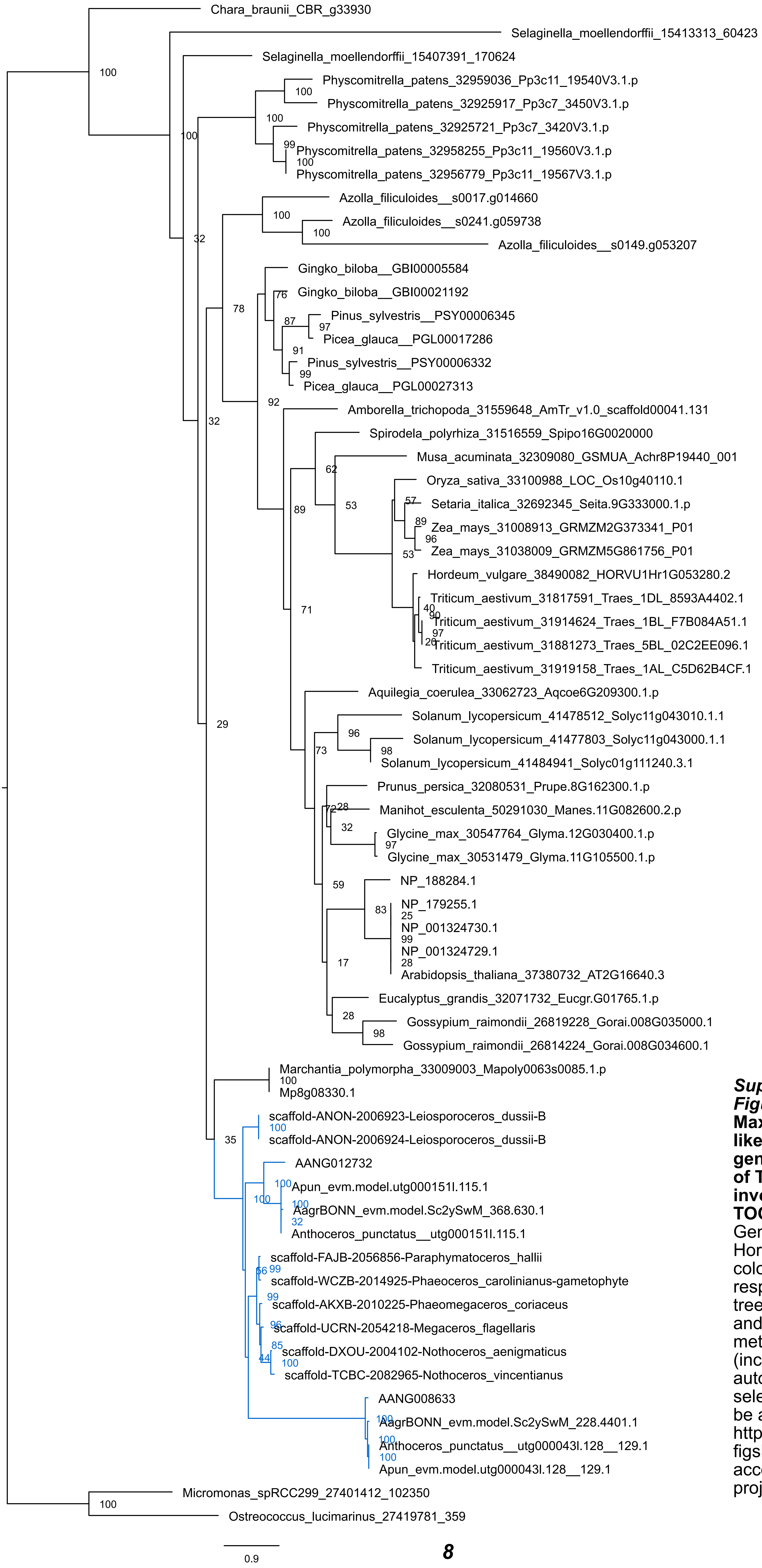


***Supplementary Figure S8.* Maximum likelihood (ML) gene phylogeny of TOC159, involved in the TOC translocon.** Genes specific to Hornworts are coloured blue, respectively. Raw tree topologies, and relevant metadata (including automatic model selection), can be accessed at https://figshare.com/account/home#/projects/125431

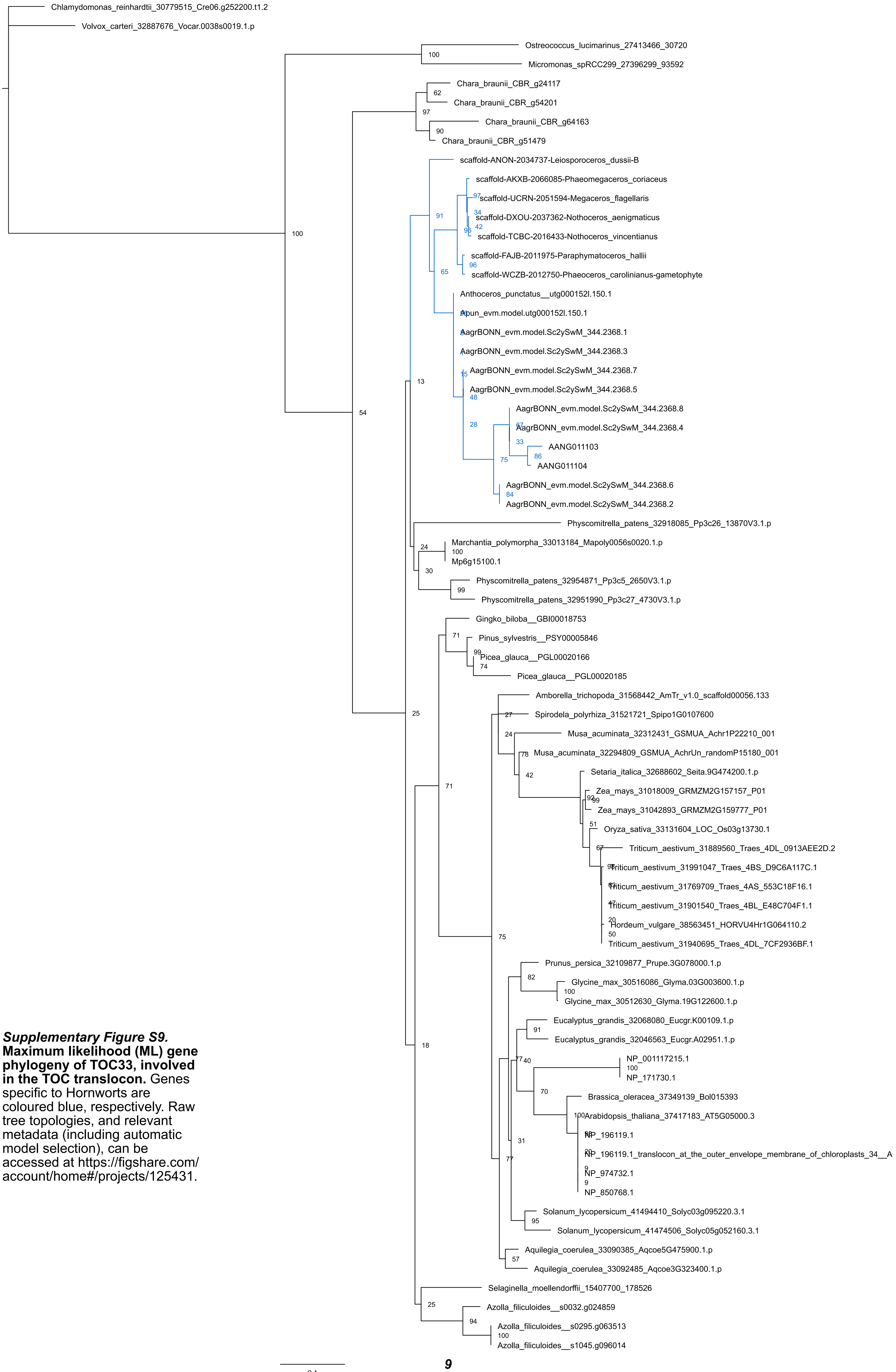


***Supplementary Figure S9.* Maximum likelihood (ML) gene phylogeny of TOC33, involved in the TOC translocon.** Genes specific to Hornworts are coloured blue, respectively. Raw tree topologies, and relevant metadata (including automatic model selection), can be accessed at https://figshare.com/account/home#/projects/125431.

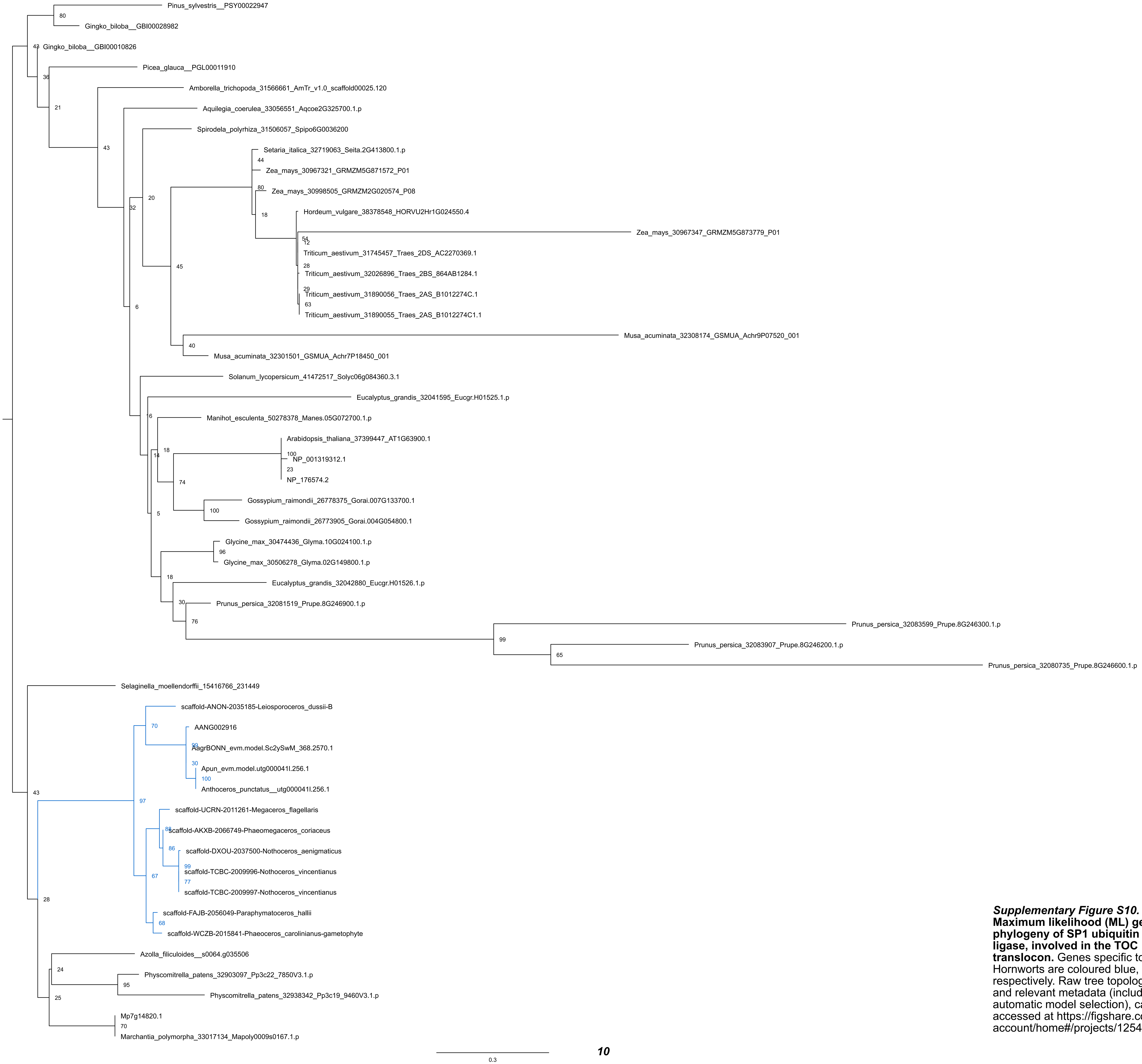


***Supplementary Figure S10.* Maximum likelihood (ML) gene phylogeny of SP1 ubiquitin ligase, involved in the TOC translocon.** Genes specific to Hornworts are coloured blue, respectively. Raw tree topologies, and relevant metadata (including automatic model selection), can be accessed at https://figshare.com/account/home#/projects/125431.

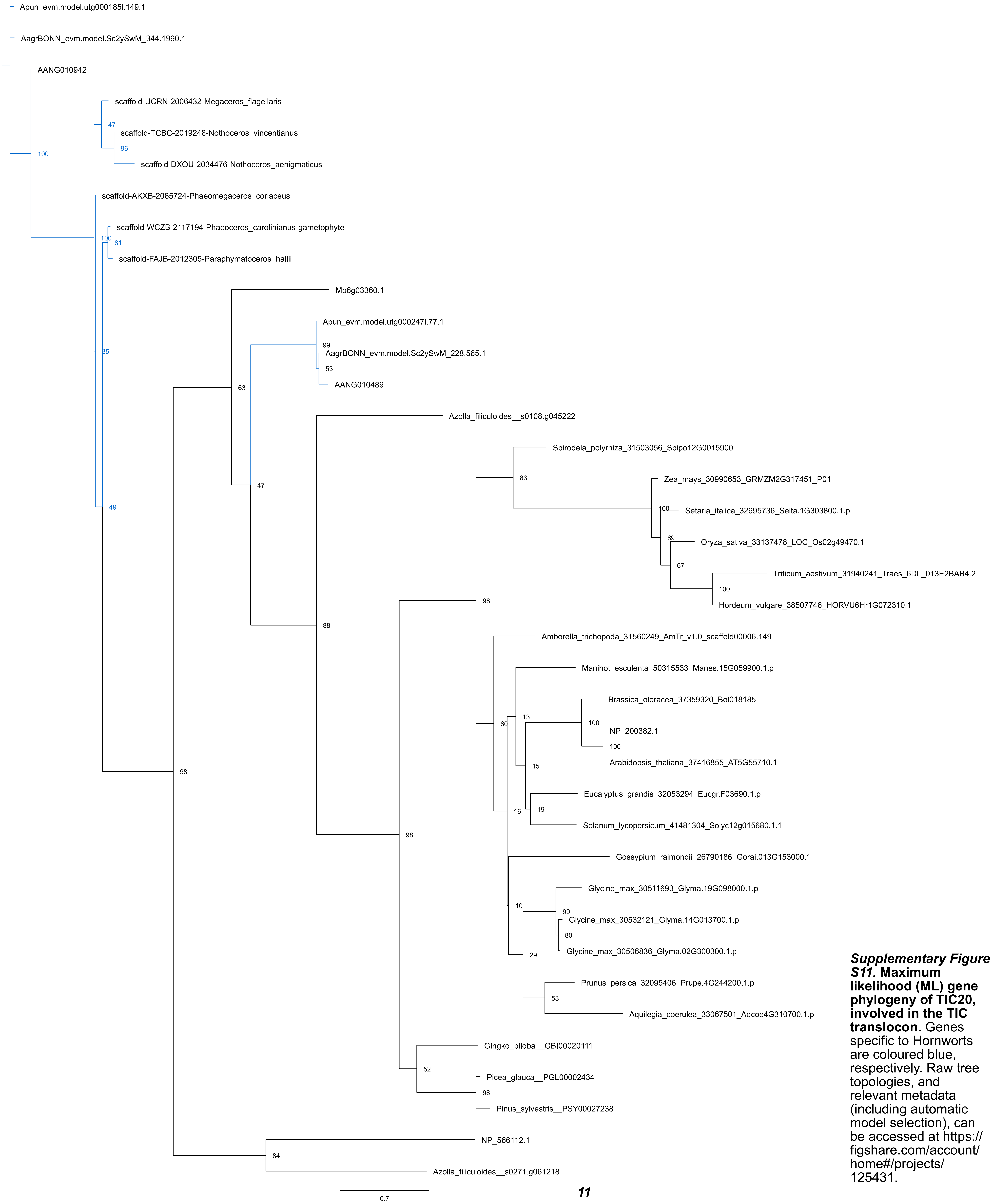


***Supplementary Figure S11.* Maximum likelihood (ML) gene phylogeny of TIC20, involved in the TIC translocon.** Genes specific to Hornworts are coloured blue, respectively. Raw tree topologies, and relevant metadata (including automatic model selection), can be accessed at https://figshare.com/account/home#/projects/125431.

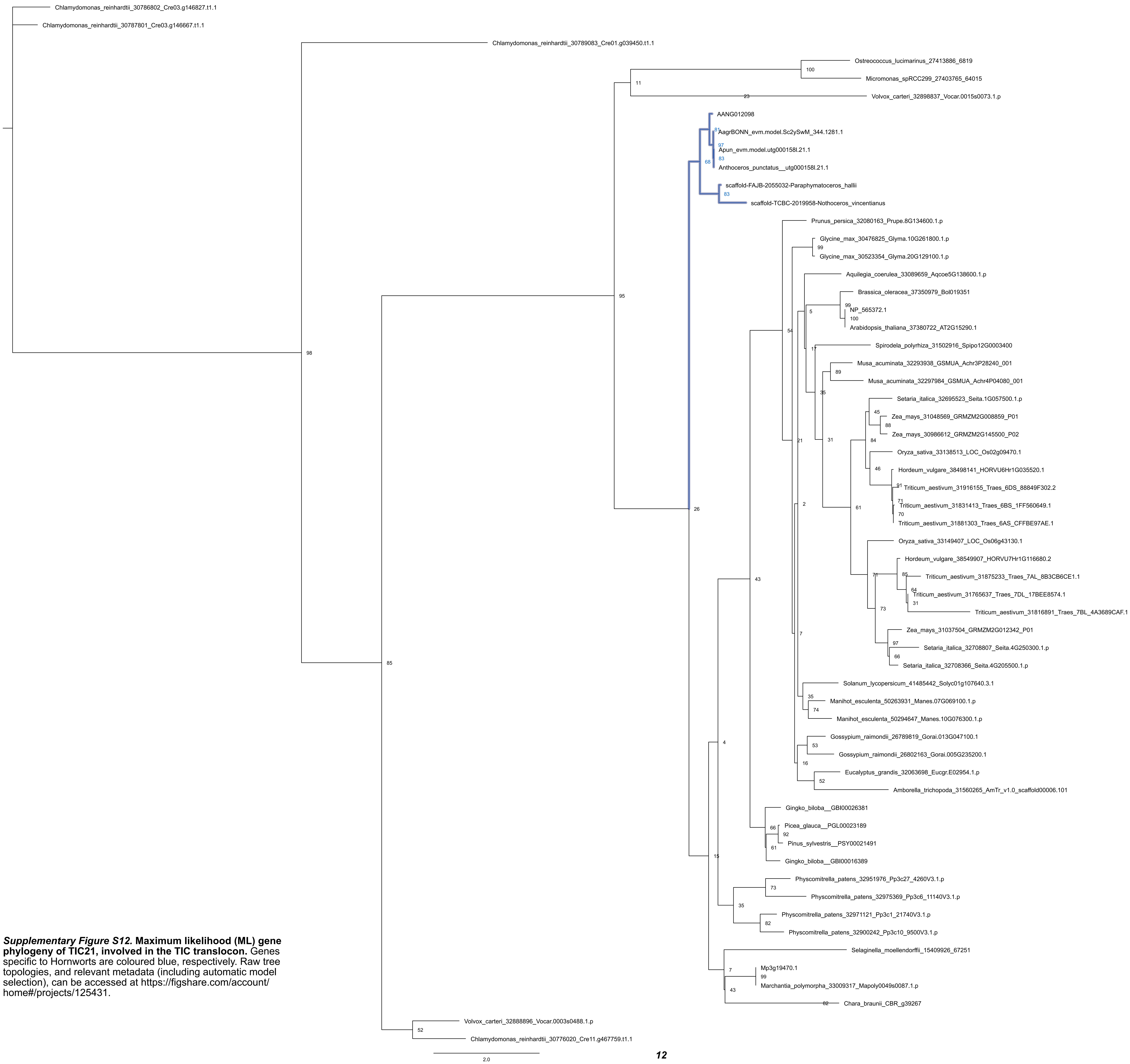


***Supplementary Figure S12.* Maximum likelihood (ML) gene phylogeny of TIC21, involved in the TIC translocon.** Genes specific to Hornworts are coloured blue, respectively. Raw tree topologies, and relevant metadata (including automatic model selection), can be accessed at https://figshare.com/account/home#/projects/125431.

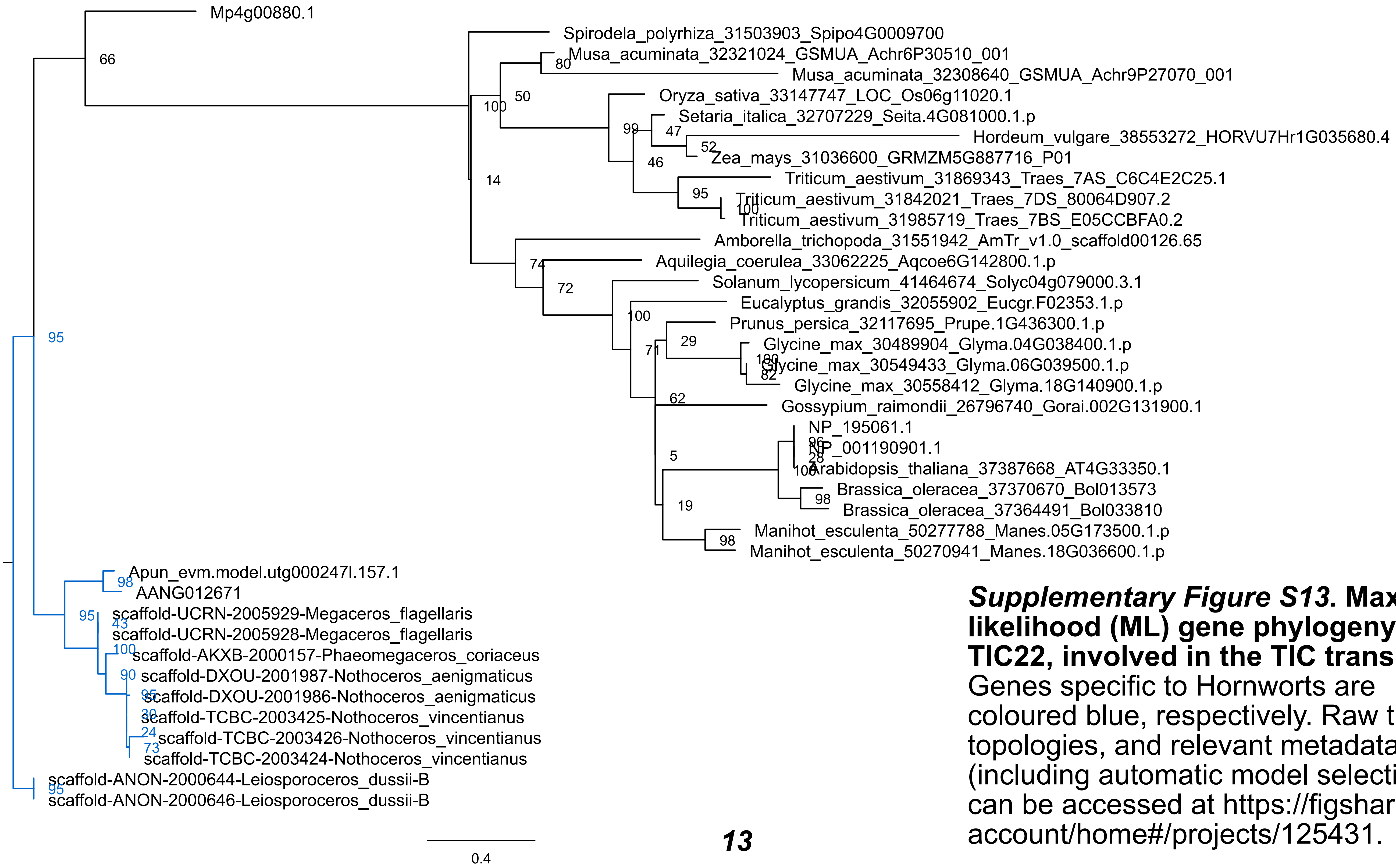


***Supplementary Figure S13.* Maximum likelihood (ML) gene phylogeny of TIC22, involved in the TIC translocon.** Genes specific to Hornworts are coloured blue, respectively. Raw tree topologies, and relevant metadata (including automatic model selection), can be accessed at https://figshare.com/account/home#/projects/125431.

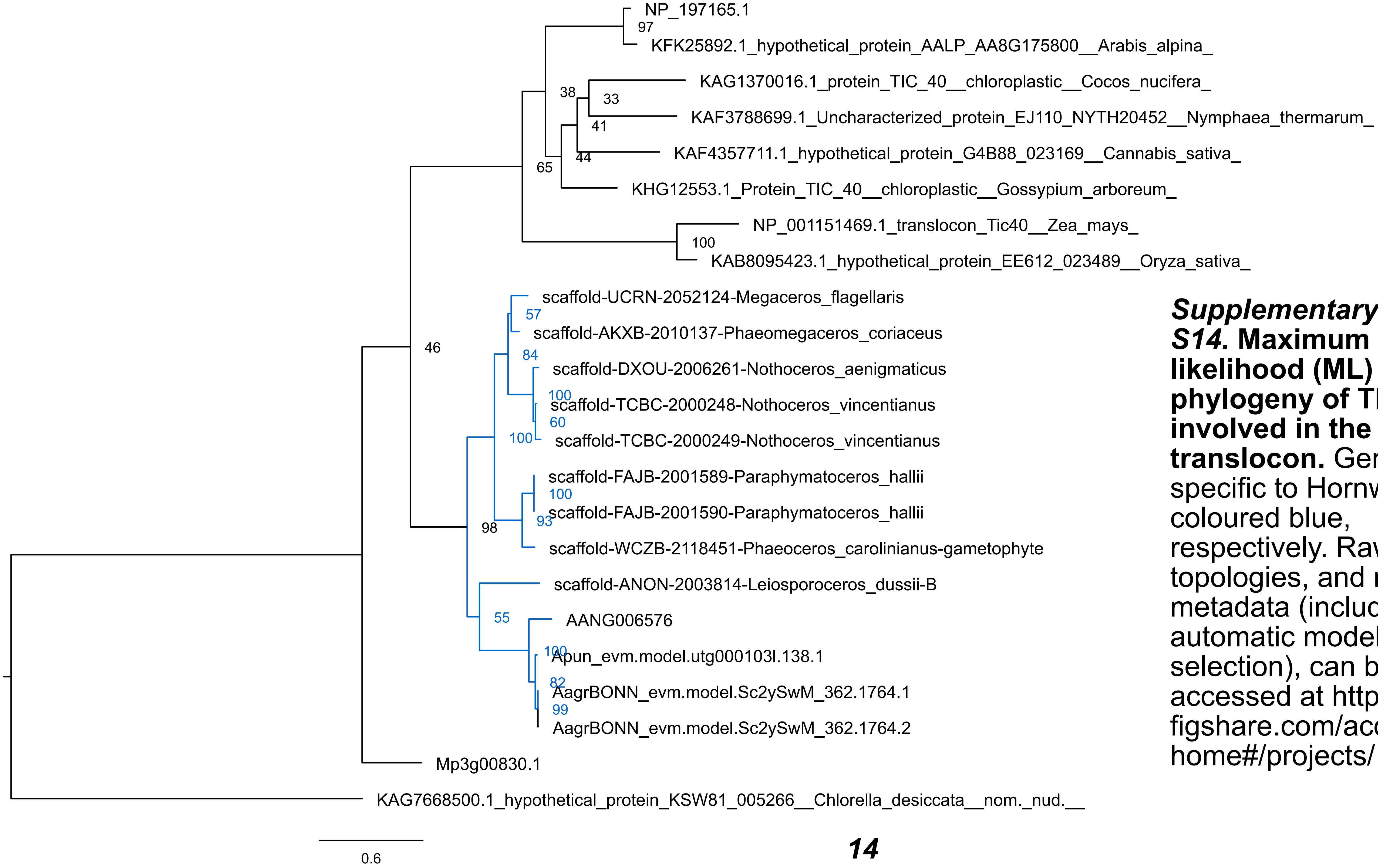


***Supplementary Figure S14.* Maximum likelihood (ML) gene phylogeny of TIC40, involved in the TIC translocon.** Genes specific to Hornworts are coloured blue, respectively. Raw tree topologies, and relevant metadata (including automatic model selection), can be accessed at https://figshare.com/account/home#/projects/125431.

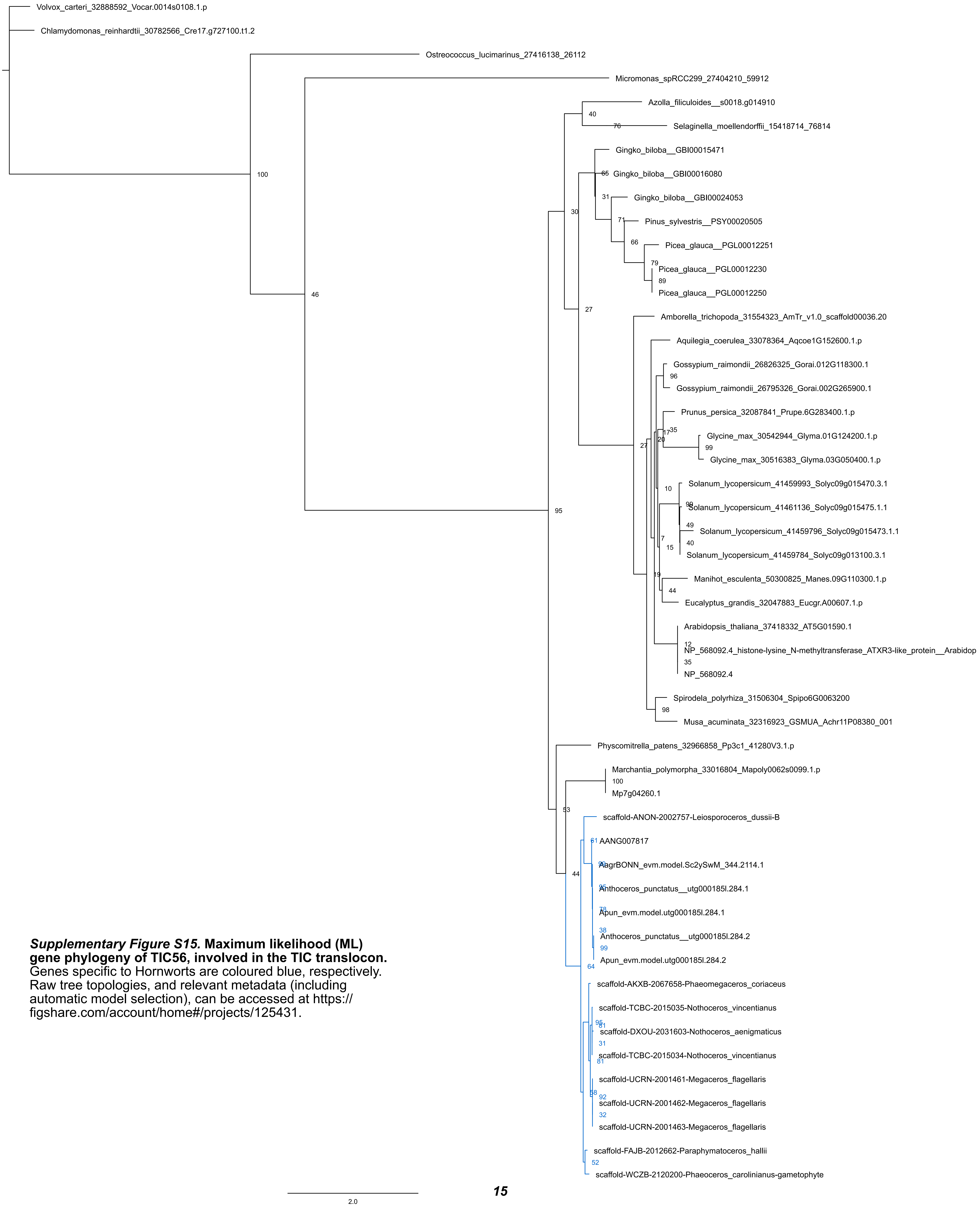


***Supplementary Figure S15.* Maximum likelihood (ML) gene phylogeny of TIC56, involved in the TIC translocon.** Genes specific to Hornworts are coloured blue, respectively. Raw tree topologies, and relevant metadata (including automatic model selection), can be accessed at https://figshare.com/account/home#/projects/125431.

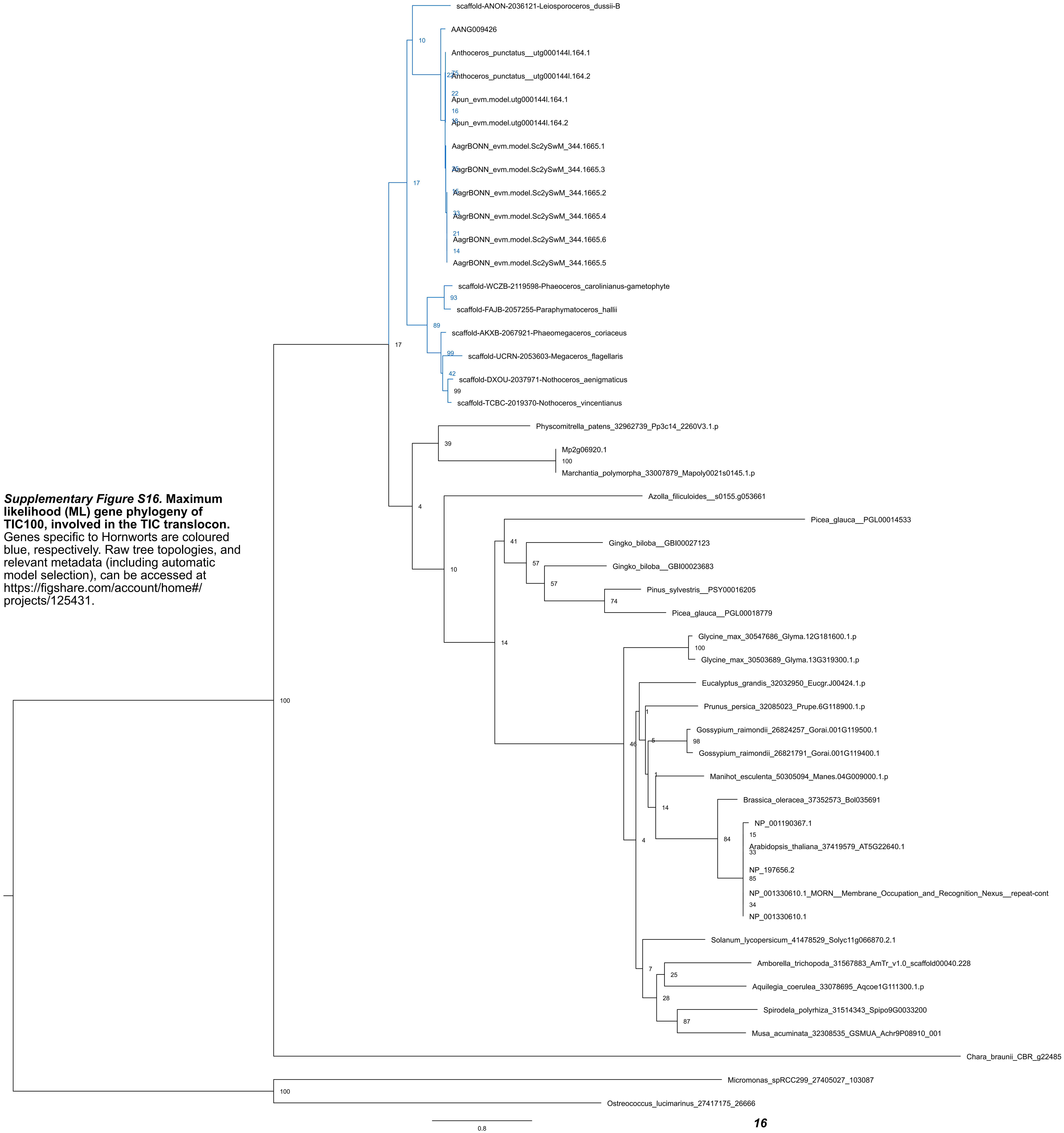


***Supplementary Figure S16.*** **Maximum likelihood (ML) gene phylogeny of TIC100, involved in the TIC translocon.** Genes specific to Hornworts are coloured blue, respectively. Raw tree topologies, and relevant metadata (including automatic model selection), can be accessed at https://figshare.com/account/home#/projects/125431.

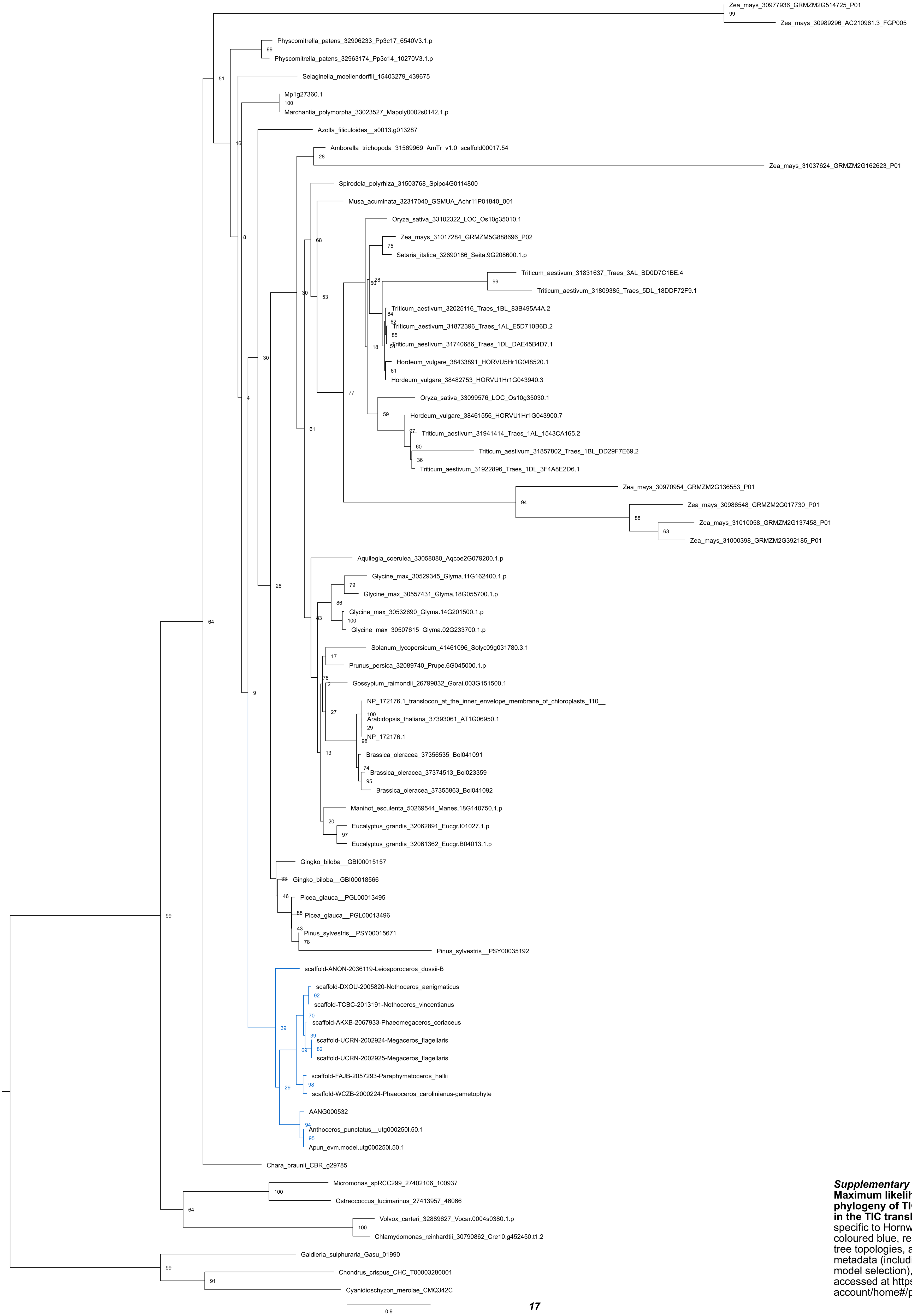


***Supplementary Figure S17.* Maximum likelihood (ML) gene phylogeny of TIC110, involved in the TIC translocon.** Genes specific to Hornworts are coloured blue, respectively. Raw tree topologies, and relevant metadata (including automatic model selection), can be accessed at https://figshare.com/account/home#/projects/125431.

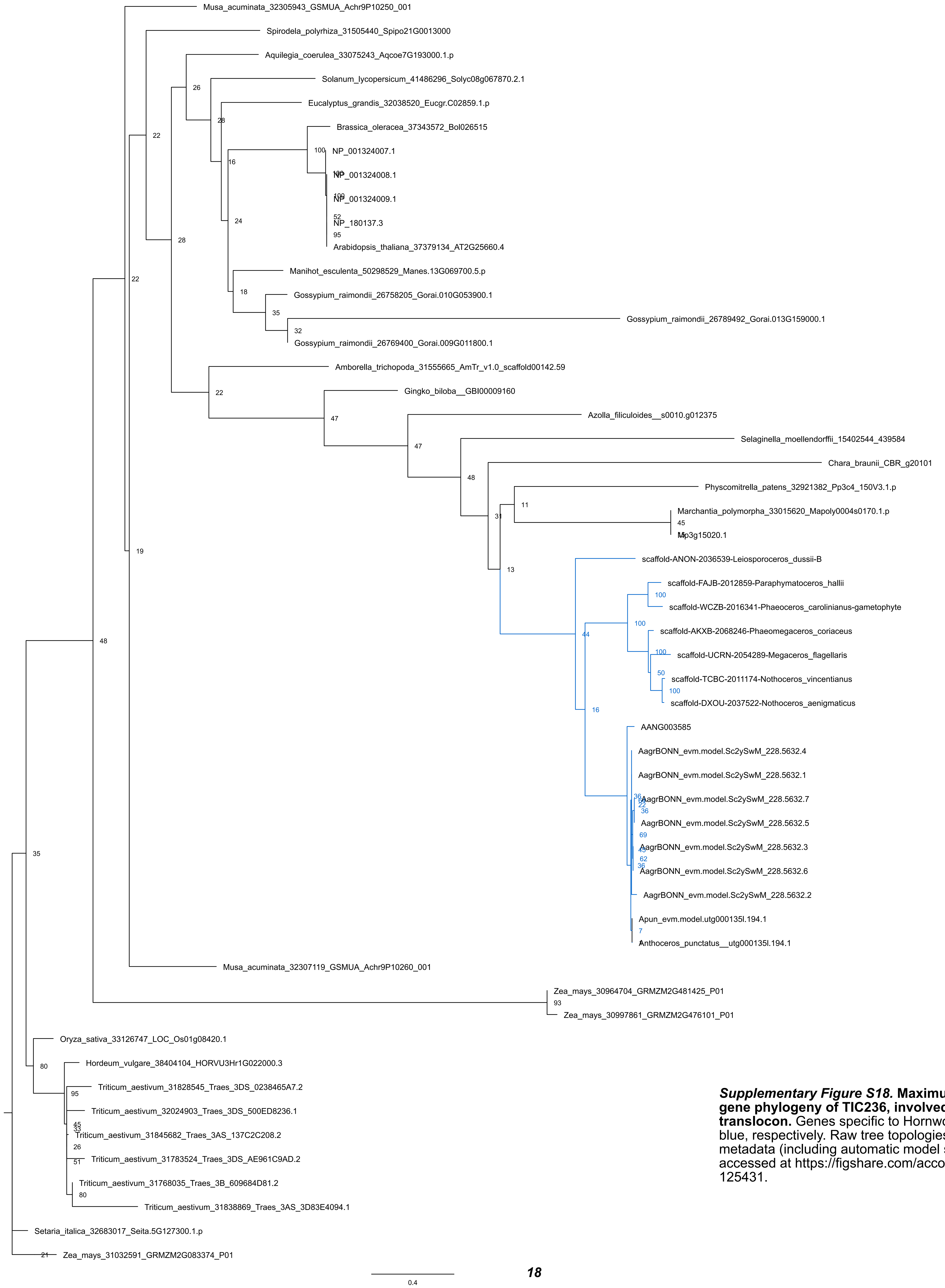


***Supplementary Figure S18.*** **Maximum likelihood (ML) gene phylogeny of TIC236, involved in the TIC translocon.** Genes specific to Hornworts are coloured blue, respectively. Raw tree topologies, and relevant metadata (including automatic model selection), can be accessed at https://figshare.com/account/home#/projects/125431.

*Supplementary Figure S19.* **Maximum likelihood (ML) gene phylogeny of TIC214/ YCF1, involved in the TIC translocon.** Genes specific to Hornworts are coloured blue, respectively. Raw tree topologies, and relevant metadata (including automatic model selection), can be accessed at https://figshare.com/account/home#/projects/125431.

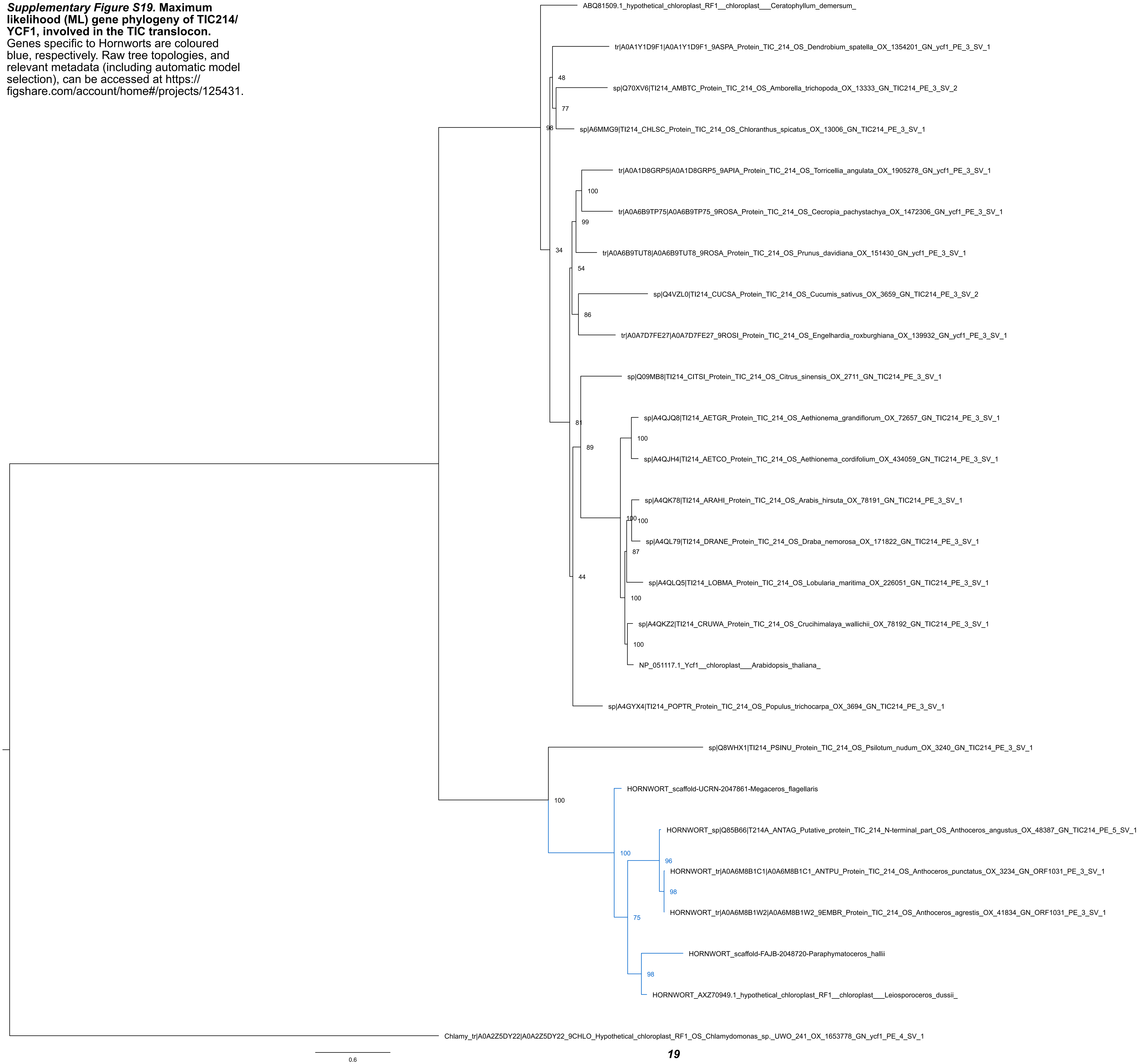

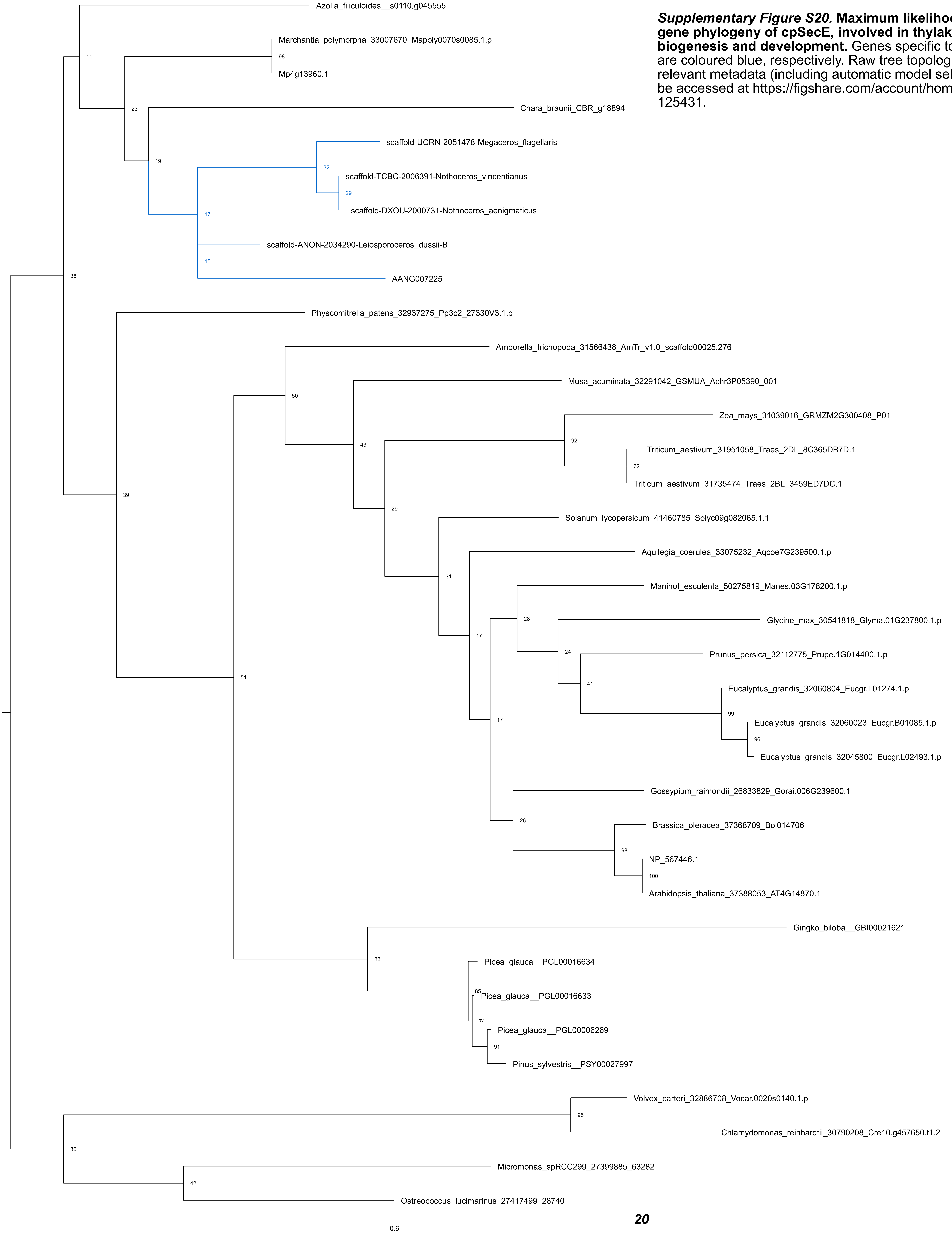


***Supplementary Figure S20.* Maximum likelihood (ML) gene phylogeny of cpSecE, involved in thylakoid biogenesis and development.** Genes specific to Hornworts are coloured blue, respectively. Raw tree topologies, and relevant metadata (including automatic model selection), can be accessed at https://figshare.com/account/home#/projects/125431.

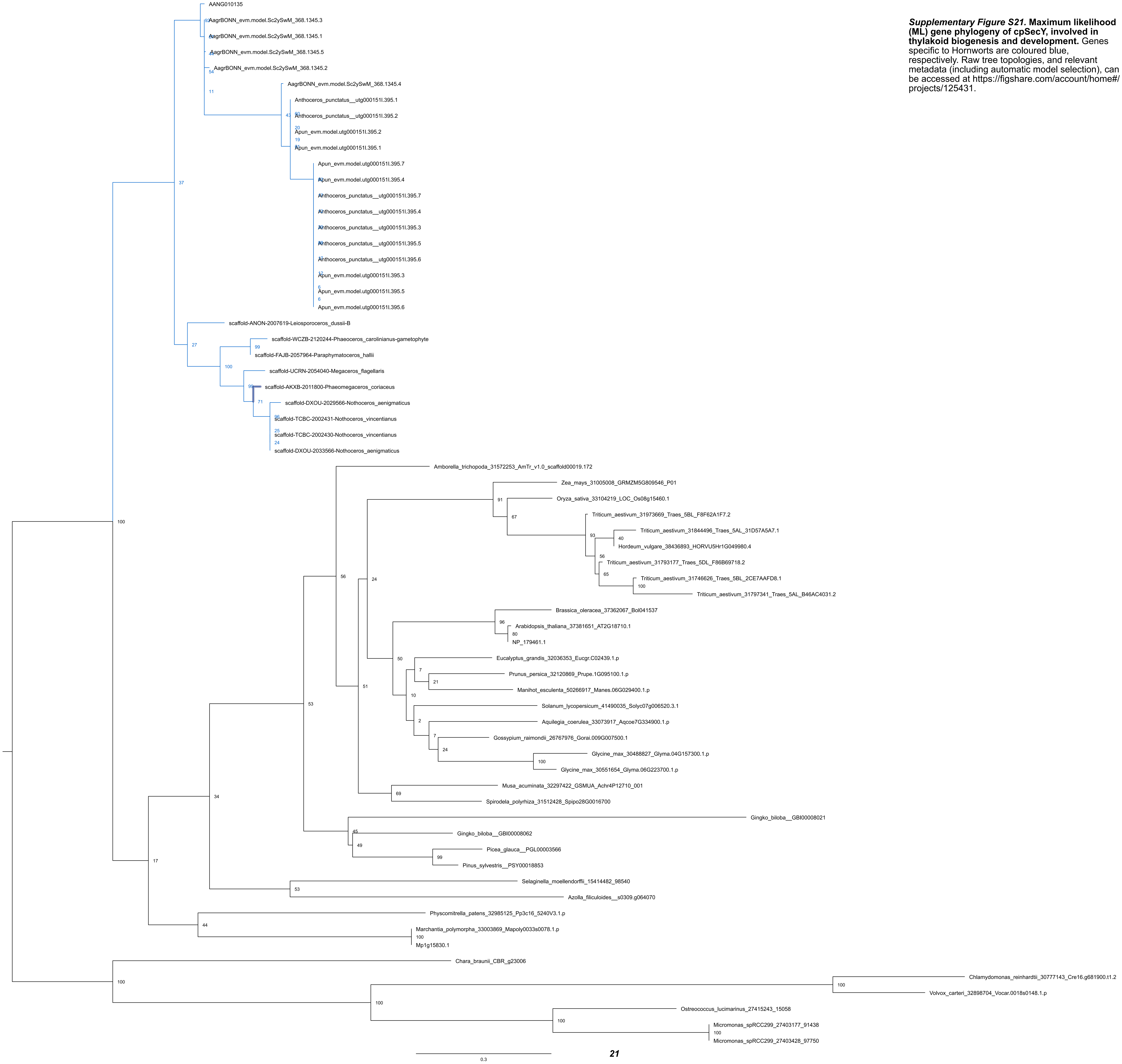

***Supplementary Figure S21.* Maximum likelihood (ML) gene phylogeny of cpSecY, involved in thylakoid biogenesis and development.** Genes specific to Hornworts are coloured blue, respectively. Raw tree topologies, and relevant metadata (including automatic model selection), can be accessed at https://figshare.com/account/home#/projects/125431.

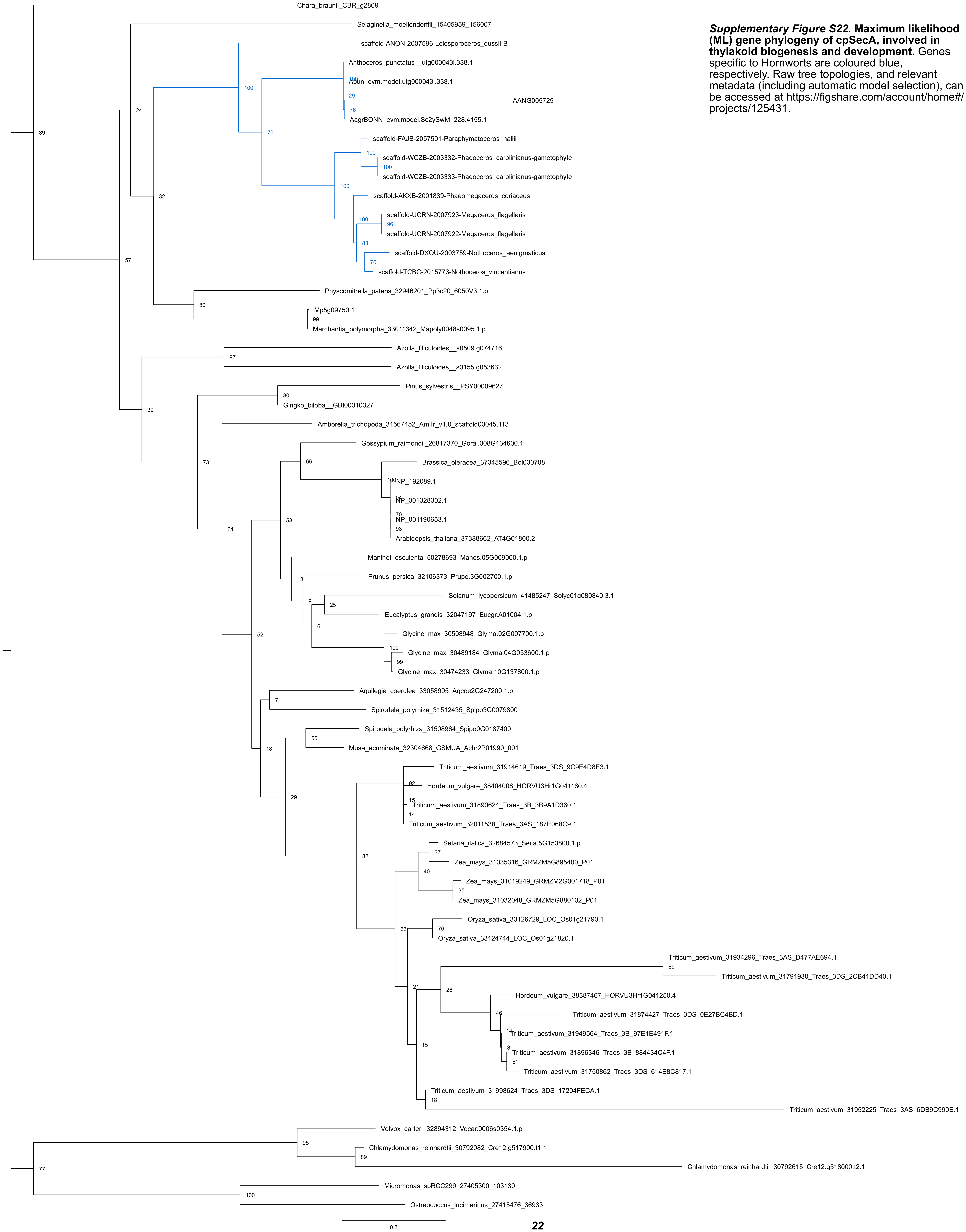


***Supplementary Figure S22.*** **Maximum likelihood (ML) gene phylogeny of cpSecA, involved in thylakoid biogenesis and development.** Genes specific to Hornworts are coloured blue, respectively. Raw tree topologies, and relevant metadata (including automatic model selection), can be accessed at https://figshare.com/account/home#/projects/125431.

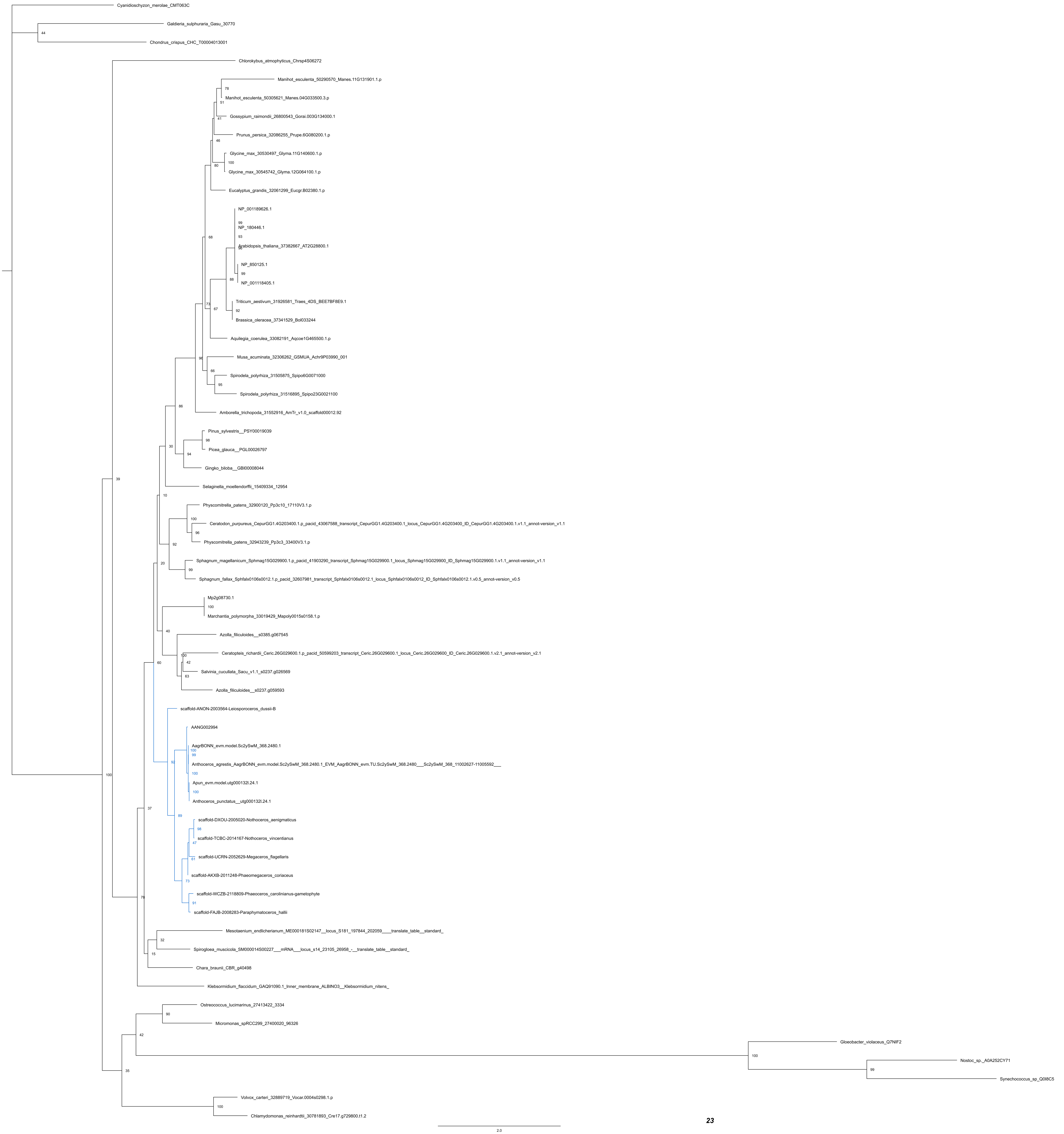


***Supplementary Figure S23.* Maximum likelihood (ML) gene phylogeny of ALB3, involved in thylakoid biogenesis and development.** Genes specific to Hornworts are coloured blue, respectively. Raw tree topologies, and relevant metadata (including automatic model selection), can be accessed at https://figshare.com/account/home#/projects/125431.

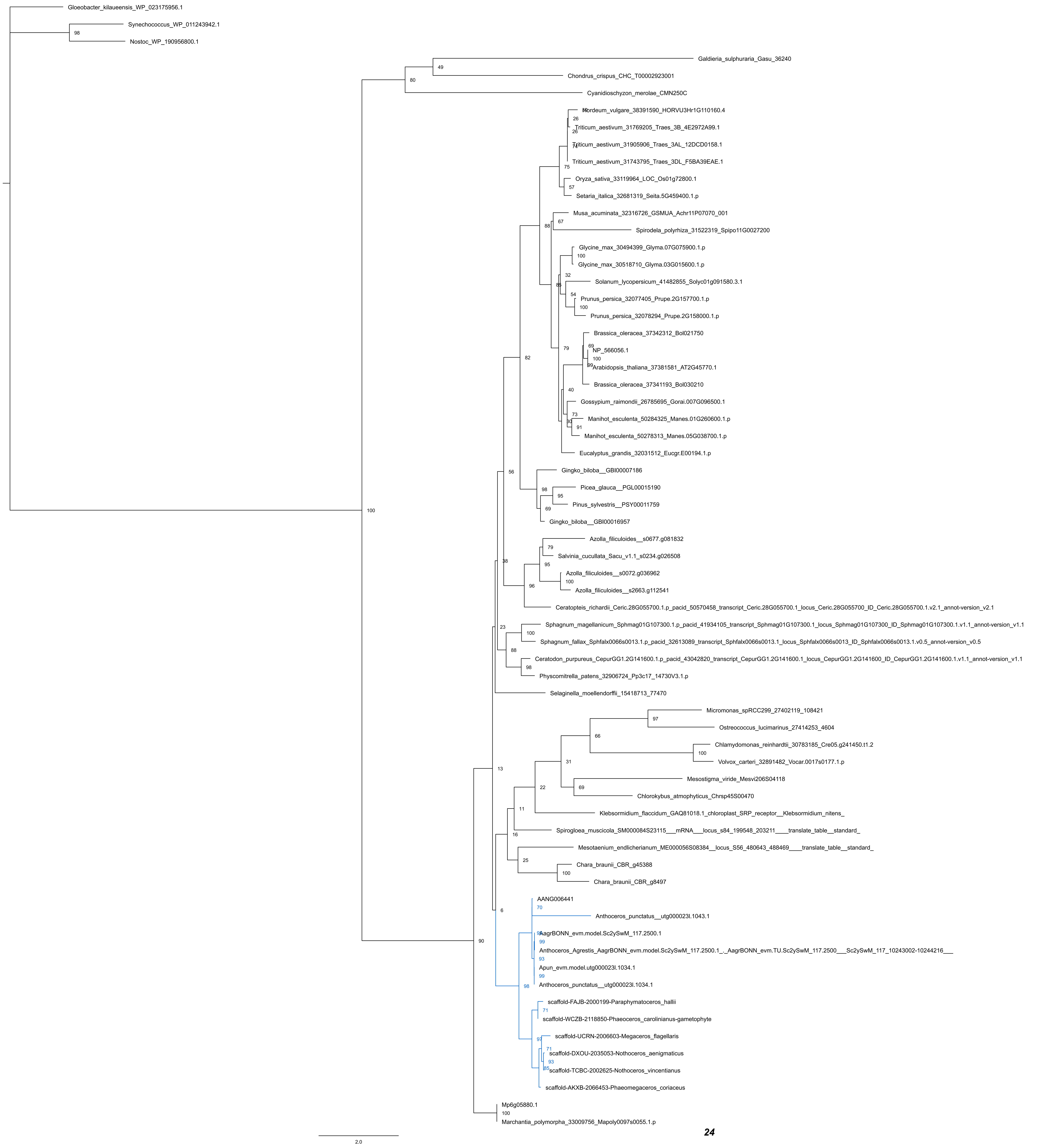

***Supplementary Figure S24.* Maximum likelihood (ML) gene phylogeny of FtsY, involved in thylakoid biogenesis and development.** Genes specific to Hornworts are coloured blue, respectively. Raw tree topologies, and relevant metadata (including automatic model selection), can be accessed at https://figshare.com/account/home#/projects/125431.

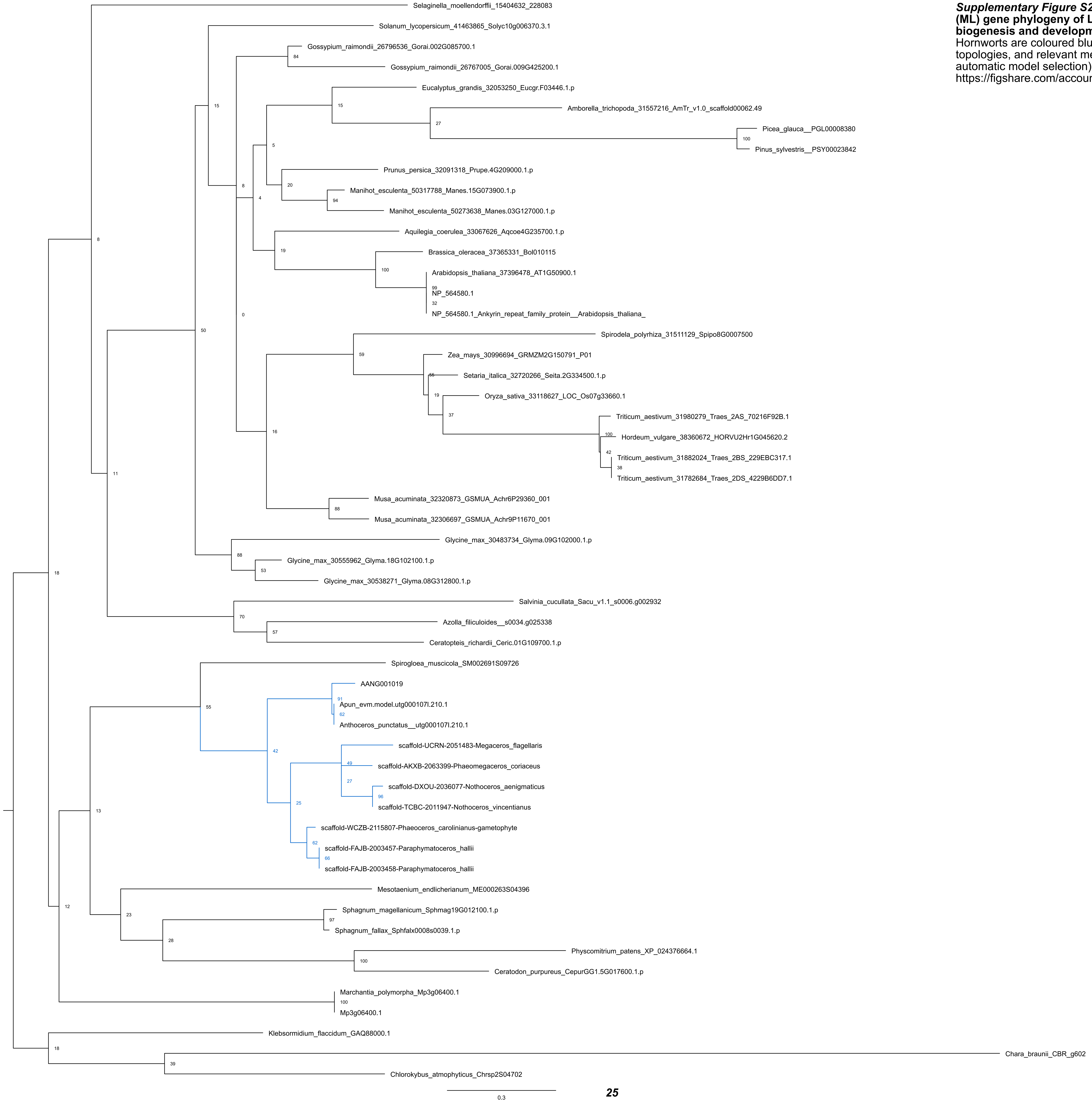


***Supplementary Figure S25.* Maximum likelihood (ML) gene phylogeny of LTD, involved in thylakoid biogenesis and development.** Genes specific to Hornworts are coloured blue, respectively. Raw tree topologies, and relevant metadata (including automatic model selection), can be accessed at https://figshare.com/account/home#/projects/125431.

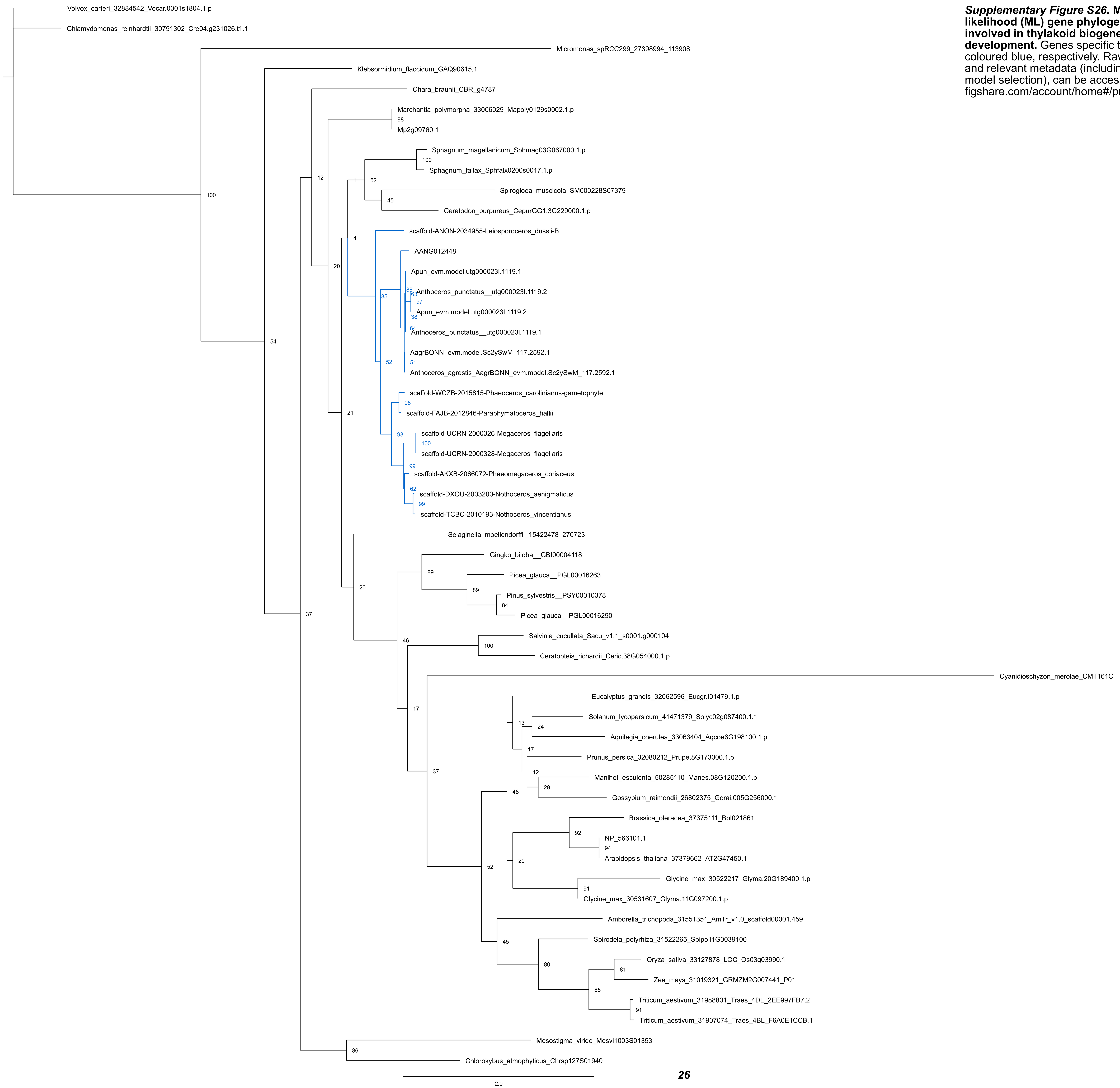


***Supplementary Figure S26.*** **Maximum likelihood (ML) gene phylogeny of SRP43, involved in thylakoid biogenesis and development.** Genes specific to Hornworts are coloured blue, respectively. Raw tree topologies, and relevant metadata (including automatic model selection), can be accessed at https://figshare.com/account/home#/projects/125431.

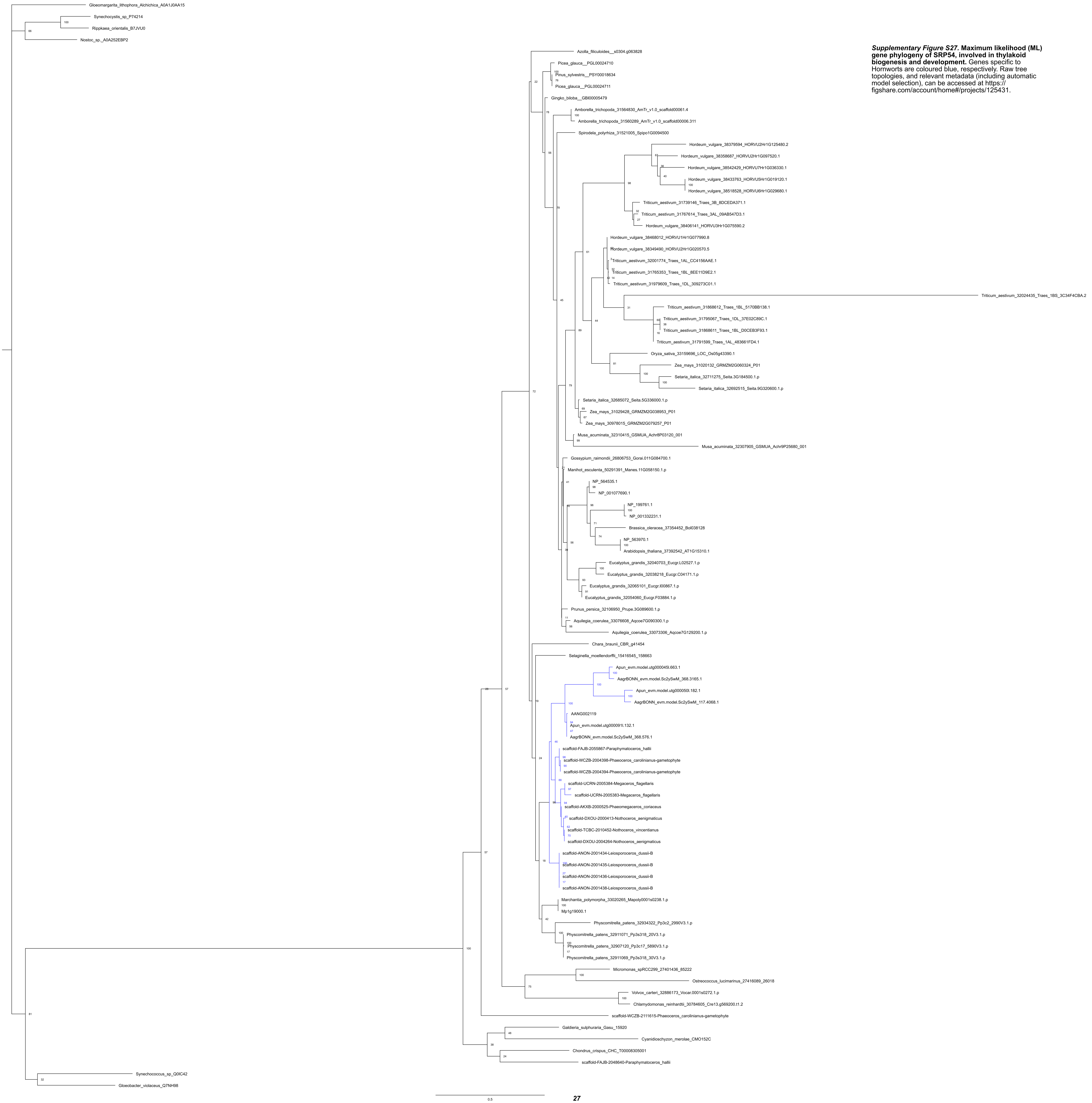


***Supplementary Figure S27*. Maximum likelihood (ML) gene phylogeny of SRP54, involved in thylakoid biogenesis and development.** Genes specific to Hornworts are coloured blue, respectively. Raw tree topologies, and relevant metadata (including automatic model selection), can be accessed at https://figshare.com/account/home#/projects/125431.

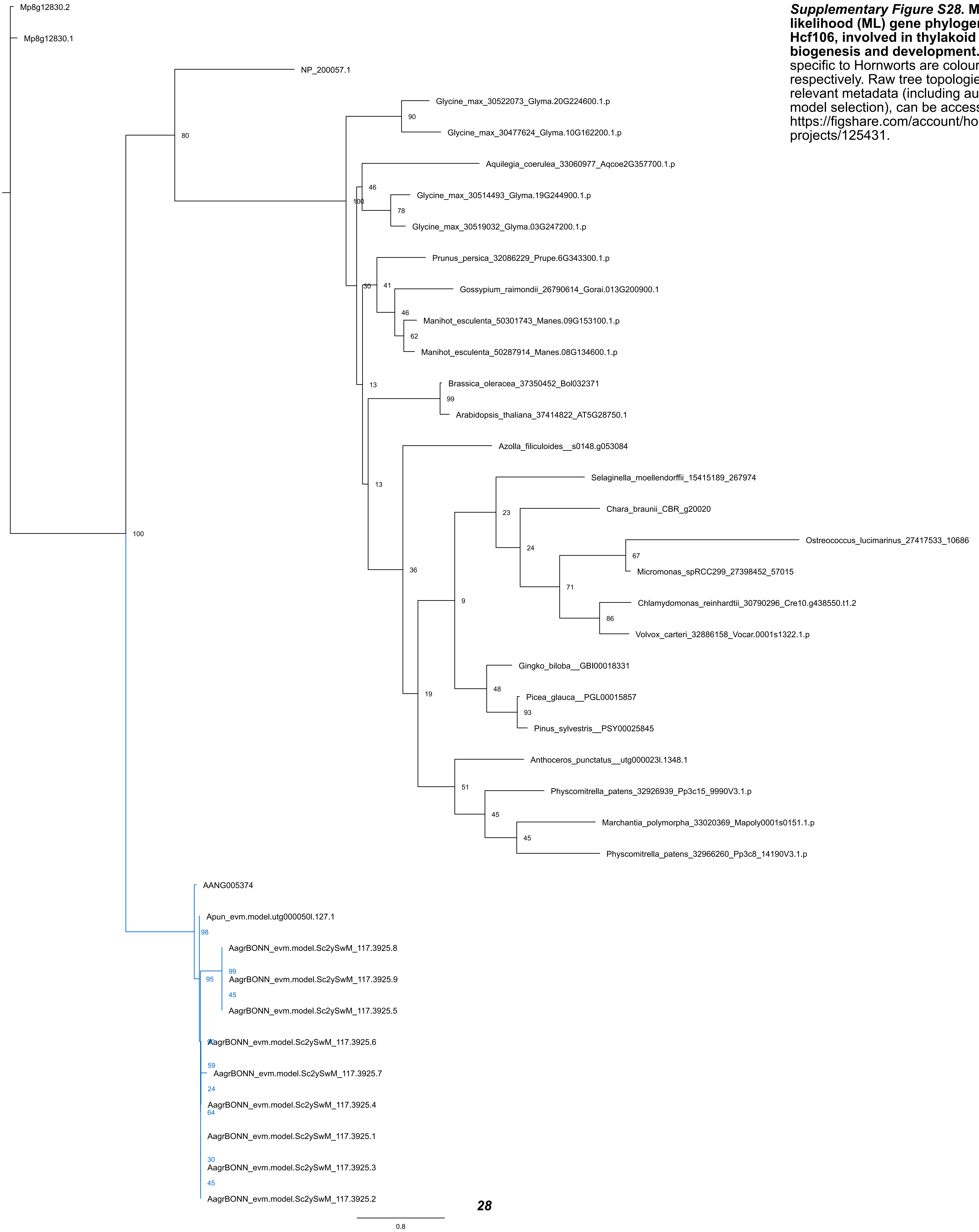


***Supplementary Figure S28.*** **Maximum likelihood (ML) gene phylogeny of Hcf106, involved in thylakoid biogenesis and development.** Genes specific to Hornworts are coloured blue, respectively. Raw tree topologies, and relevant metadata (including automatic model selection), can be accessed at https://figshare.com/account/home#/projects/125431.

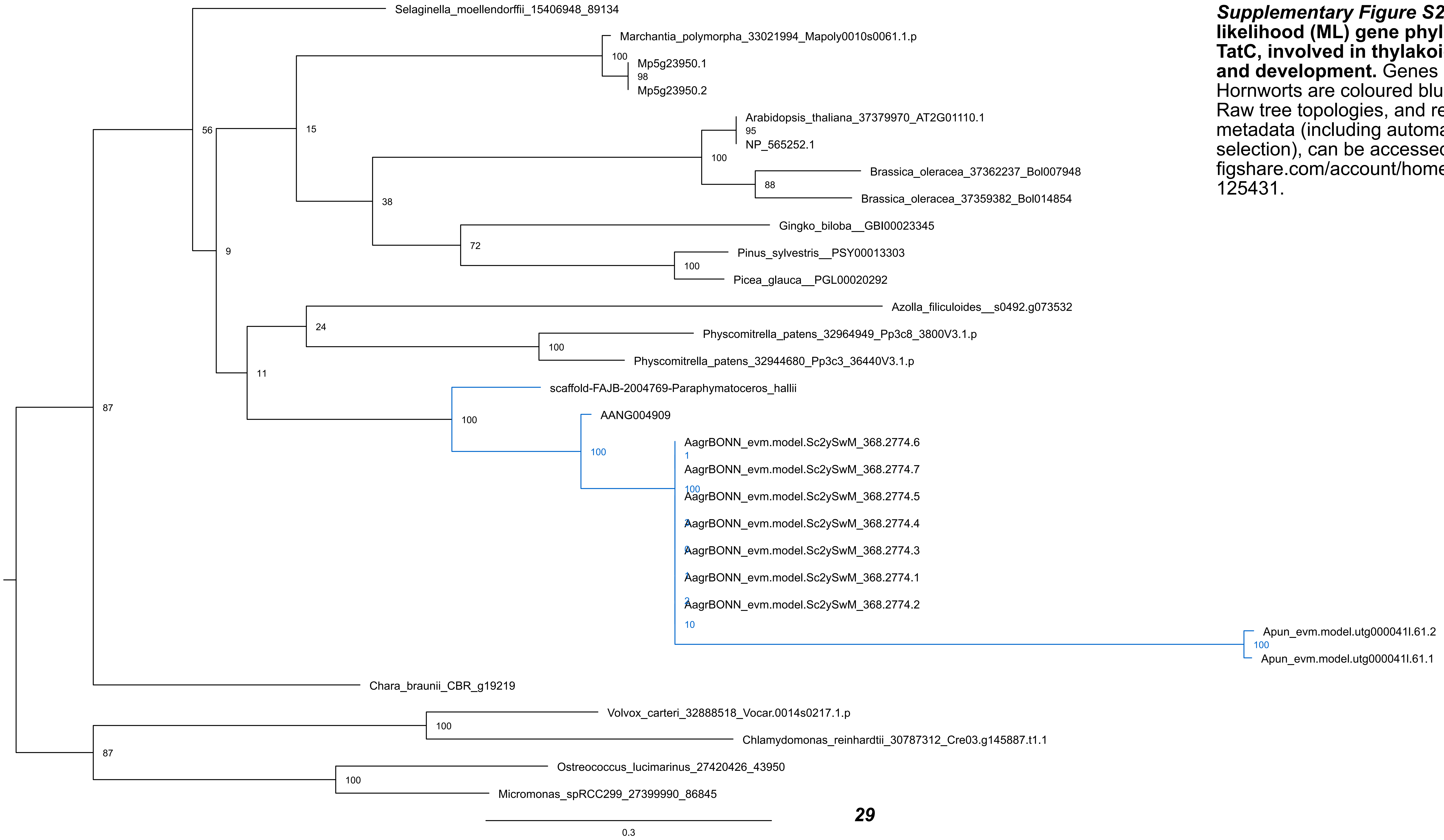


***Supplementary Figure S29.* Maximum likelihood (ML) gene phylogeny of TatC, involved in thylakoid biogenesis and development.** Genes specific to Hornworts are coloured blue, respectively. Raw tree topologies, and relevant metadata (including automatic model selection), can be accessed at https://figshare.com/account/home#/projects/125431.

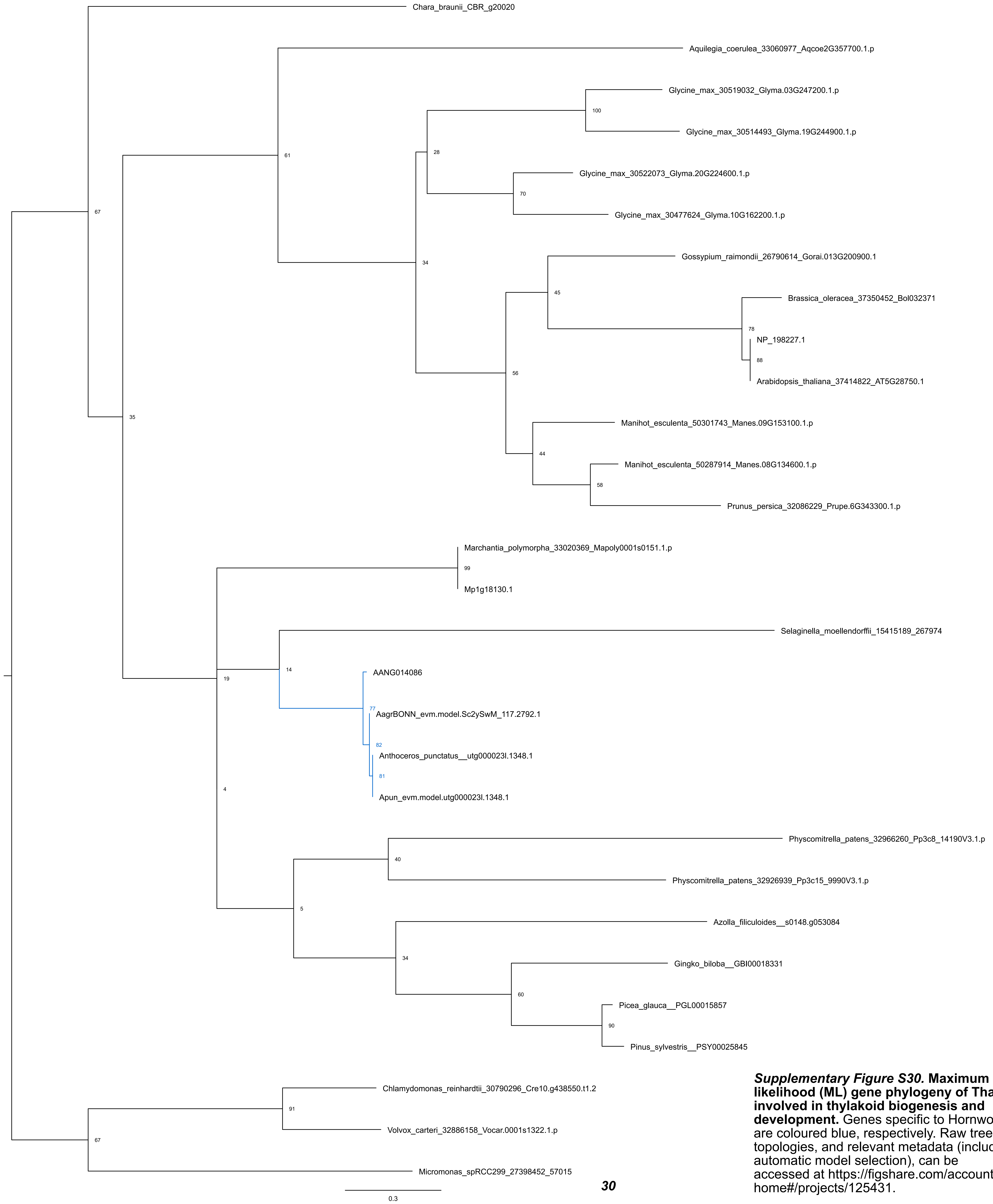


***Supplementary Figure S30.*** **Maximum likelihood (ML) gene phylogeny of Tha4, involved in thylakoid biogenesis and development.** Genes specific to Hornworts are coloured blue, respectively. Raw tree topologies, and relevant metadata (including automatic model selection), can be accessed at https://figshare.com/account/home#/projects/125431.

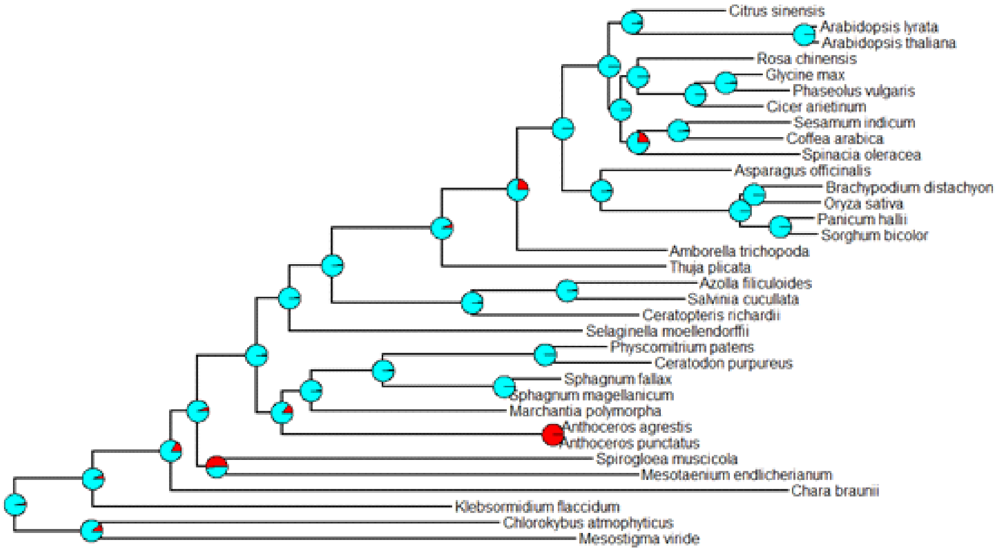


***Supplementary Figure S31***. ASR analysis showing the probability for the presence/ absence of FtsZ2 at various nodes in the tree. Presence is indicated by cyan; absence by red.

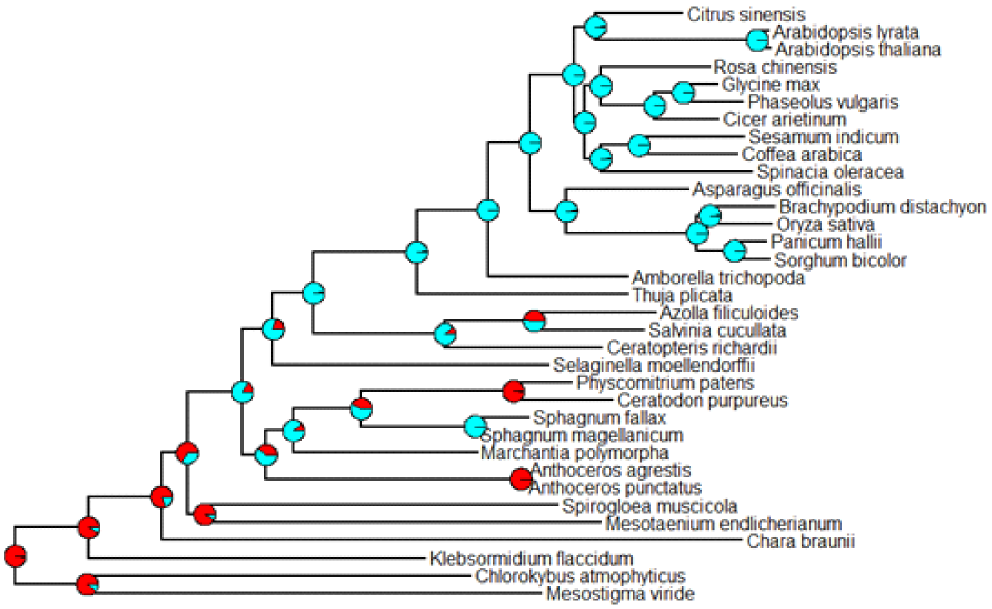


***Supplementary Figure S32***. ASR analysis showing the probability for the presence/ absence of ARC3 at various nodes in the tree. Presence is indicated by cyan; absence by red.
